# *Arhgef18* is a component of the outer limiting membrane required for retinal homeostasis

**DOI:** 10.1101/2025.10.07.680916

**Authors:** Ana Alonso-Carriazo Fernandez, Tiansheng Liu, Holly Thomas, Xiaochen Fan, Karl Matter, Maria S. Balda

**Affiliations:** Institute of Ophthalmology, University College London; Bath Street, London, EC1V 9EL, UK

**Author notes:** Corresponding Authors, Address for Communications: Institute of Ophthalmology, University College London, 11-43 Bath Street, London EC1V 9EL, UK, /6861. These authors contributed equally (Ana Alonso-Carriazo Fernandez, Tiansheng Liu, Holly Thomas).

## Abstract

Biallelic *ARHGEF18* mutations cause human adult-onset retinal degeneration. We now find that Arhgef18 associates with the retinal outer limiting membrane (OLM), an adherens junction between Müller glia and photoreceptors. *Arhgef18* knockout in Müller glial cells led to OLM disruption and vision loss by P60. While mice developed morphologically normal retinas, retinal rosettes started to form by P8, and retinas then progressively degenerated with OLM disintegration, retinal thinning, and vascular leakage. ARHGEF18/p114RhoGEF depletion in Müller cells in culture confirmed disruption of junctional recruitment of OLM proteins. Depletion also induced activation of NF-κB and β-catenin signalling, activation of the multifunctional kinase Tank-binding kinase 1 (TBK1) and reduced mitochondrial activity. TBK1 inhibition or directly supporting mitochondrial activity with nicotinamide attenuated NF-κB and β-catenin signalling and rescued mitochondrial activity. Thus, Arhgef18 is essential for OLM maintenance, and its disruption leads to activation of mechanisms that are targetable for possible therapeutic approaches.

## Main

Biallelic *ARHGEF18* mutations that partially inactivate the protein cause adult-onset retinal degeneration, indicating that *ARHGEF18* is important for retinal homeostasis^1^. ARHGEF18 encodes p114RhoGEF, a guanine nucleotide exchange factor for RhoA that regulates the actin cytoskeleton and apicobasal polarisation^2^. Spatially restricted activation of RhoA at cell-cell contacts by p114RhoGEF drives epithelial and endothelial junction formation and morphogenesis, and is required for junctional integrity under mechanical stress^2–4^. Inactivation of ArhGEF18 in the medaka fish increases proliferation of retinal progenitors and leads to larval lethality^5^. In mice, global *Arhgef18* knockout is embryonic lethal^6^. Thus, to investigate the role of ARHGEF18 in the retina and how its malfunction causes retinal disease requires cell-specific mouse knockout models.

Retinal degeneration with biallelic mutations of *ARHGEF18* resembles the one of patients with *CRB1* mutations, which induces retinitis pigmentosa (RP) and intraretinal cysts^1^. Mouse knockout of *Crb1* and *Crb2* in retinal progenitors or Müller glial cells (MGCs using a *Pdgfra-Cre* allele) leads to retinitis pigmentosa or Leber congenital amaurosis phenotypes, with defects in the outer limiting membrane^7^. MGCs are the main macroglia of the neural retina, expanding radially across all retinal layers and forming an essential scaffold for the orientation and support of retinal neurons^8^. The soma of MGCs resides in the inner nuclear layer (INL), and their processes expand from the outer nuclear layer (ONL) to the ganglion cell layer (GCL). At their apical end, Müller glia interact with photoreceptors, forming a specialised adherens junction, the outer limiting membrane (OLM)^9–11^. The OLM contains components found in other cell types in tight and adherens junctions, such as ZO-1, N-cadherin, and catenins, as well as polarity proteins, such as CRB1, CRB2, PALS1 and MUPP1^12^.

As ARHGEF18/p114RhoGEF plays a role in cell-cell adhesion complexes, we hypothesised that it functions at the OLM, supporting adhesion and retinal integrity. Hence, we employed the *Pdgfra-Cre* allele to knockout *Arhgef18* in MGCs. We found that Arhgef18/p114RhoGEF is indeed a component of the OLM. *Arhgef18* knockout in MGCs induced early onset retinal degeneration, starting with formation of retinal rosettes at P8 and disruption of cell-cell adhesion at the OLM, and led to a progressive loss of retinal structure with vascular leakage and vision loss. These results demonstrate that Arhgef18 is a functional component of the OLM required for retinal homeostasis. ARHGEF18/p114RhoGEF depletion in an MGC cell line by RNA interference led to the disruption of adherens junctions and activation of TBK1, NF-κΒ and β-catenin signalling, as well as reduction in mitochondrial respiration, indicating corrupted energy metabolism. *In vivo*, mitochondrial localisation was disrupted in the adhesion partner of MGCs, photoreceptors, upon disruption of the OLM in *Arhgef18fl/fl Pdgfra-Cre* mice. Depletion-induced responses in human MGCs could be rescued with TBK1 inhibitors or by reinforcing metabolism with nicotinamide, suggesting possible avenues to be explored as futrue therapeutic approaches.

## Results

### Arhgef18 associates with the OLM and depletion in MGCs impairs visual function

We first assessed the retinal localisation of Arhgef18/p114RhoGEF in control mice at P21. Arhgef18/p114RhoGEF was found concentrated at the OLM, where it co-localized with the established OLM components ZO-1 and p120 catenin (Fig. 1A). Some staining was observed across the retina, including the outer and inner plexiform layers, and blood vessels. Thus, Arhgef18/p114RhoGEF is a component of the OLM, and adherens junction formed by MGCs and photoreceptors.

**Figure 1:**
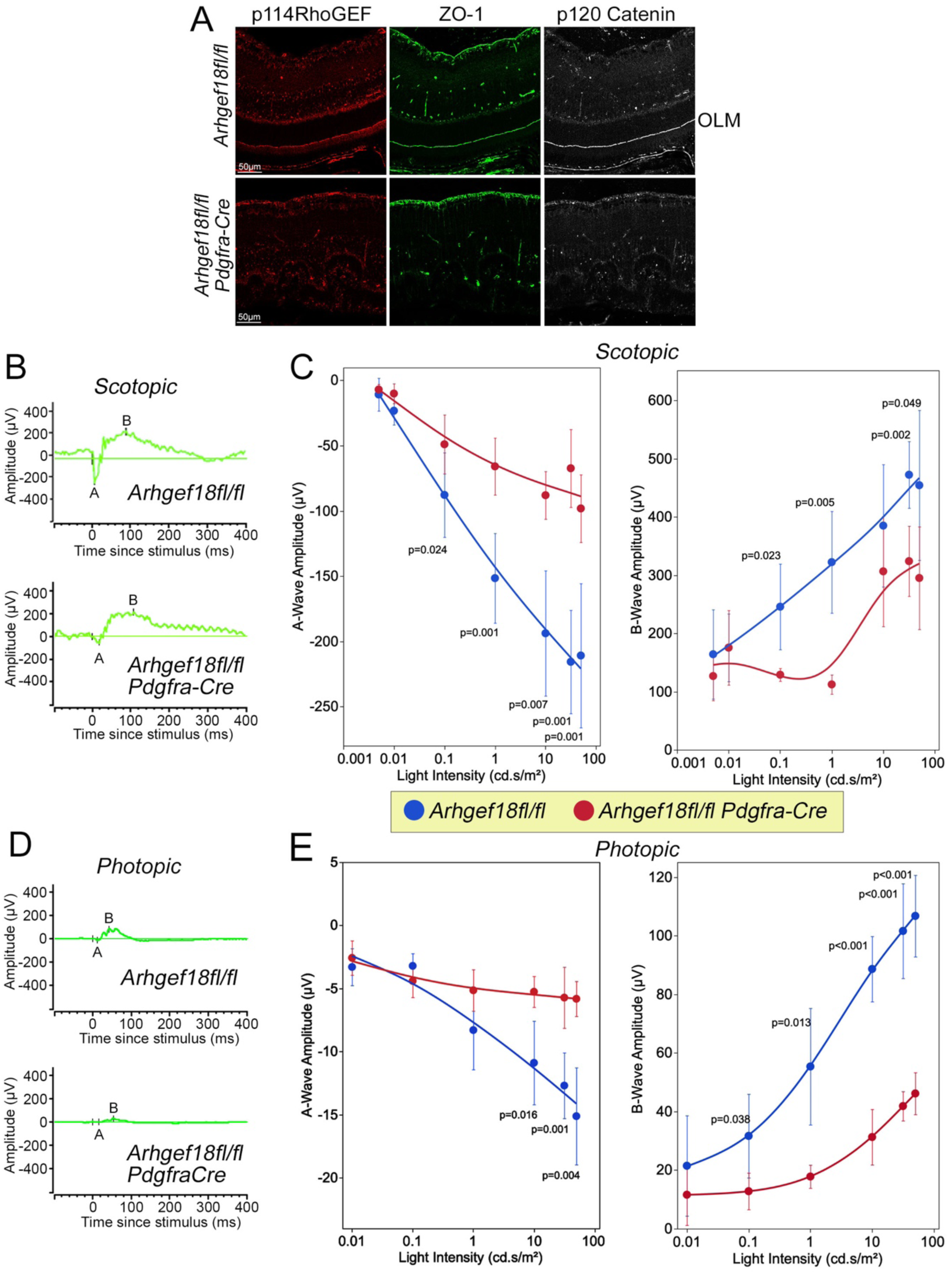
p114RhoGEF expression in MGCs is required for OLM maintenance and normal visual function. (A) Immunofluorescence of central retina sections from *Arhgef18fl/fl* and *Arhgef18fl/fl Pdgfra-Cre* mice at P21 stained for ARHGEF18/p114RhoGEF, ZO-1 and p120 Catenin. (OLM, outer limiting membrane). (B-E) Scotopic and photopic electroretinograms of 6-week-old, dark-adapted control and *Arhgef18fl/fl Pdgfra-Cre* mice at P45 (n=3). Panels B and D show example traces. Panels C and D show quantifications of A- and B-wave amplitudes (*Arhgef18fl/fl*, n=5; *Arhgef18fl/fl Pdgfra-Cre* n=7 animals). Shown are means ±1SD; p-values, two-sided t-tests comparing the two genotypes at each light intensity).

The retinal structure in patients with *ARHGEF18* biallelic mutations resembled to the one of subjects with *CRB1* mutations^1^. Mouse knockout of *Crb1* and *Crb2* in retinal progenitors or MGCs lead to defects in the OLM^7^. To investigate the role of Arhgef18/p114RhoGEF in the OLM, we therefore developed a mouse model analogous to the one described by Quinn *et al.*^7^, using a *Pdgfra-Cre* allele to knockout *Arhgef18* in MGCs. *Pdgfra-Cre* did not have deleterious effects on the retinal structure in a C57BL/6J background (Fig.S1A). However, initial analysis of *Arhgef18fl/fl Pdgfra*-*Cre* mice revealed a striking retinal degeneration with a completely disrupted OLM (Fig.1A). Arhgef18/p114RhoGEF was still apparently normally expressed in other parts of the retina, such as blood vessels and neurons of the inner retina.

Next, we examined whether *Arhgef18fl/fl Pdgfra*-*Cre* mice had visual defects. Retinal function was assessed by recording electroretinograms (ERGs) in dark-adapted mice. In 6-week-old mice, both scotopic and photopic responses were strongly reduced in *Arhgef18fl/fl Pdgfra*-*Cre* mice with A-wave amplitudes reduced by about two thirds in both scotopic and photopic ERGs, indicating defects in photoreceptor function (Fig.1B-E). Consequently, B-wave responses, measuring activity of the inner retina, were also strongly reduced. *Arhgef18* knockout in MGCs led to increased implicit times of scotopic A-waves (time period between light stimulus and A-wave minimum), a common feature of retinal degenerations affecting photoreceptors (Fig.S1B)^13^. *Pdgfra*-*Cre* mice did not exhibit any alterations in their ERG responses (Fig.S1C). Thus, *Arhgef18fl/fl Pdgfra*-*Cre* mice develop a severe retinal degeneration and a striking loss of visual function affecting rod and cone photoreceptors.

We next employed SD-OCT, fundoscopy, and fluorescent angiography (FFA) to assess how the retinal degeneration develops. SD-OCT scans showed reduced retinal thickness at P21 (∼15% reduction; Fig 2A-B) with a progressive loss of retinal thickness at week 8 and of more than 50% by week 32, with almost complete loss of identifiable retinal layers and hyperreflective scars across the remaining retina (Fig.S2). Thus, *Arhgef18* knockout in MGCs causes ONL disruption, retinal degeneration, and photoreceptor defects.

**Figure 2:**
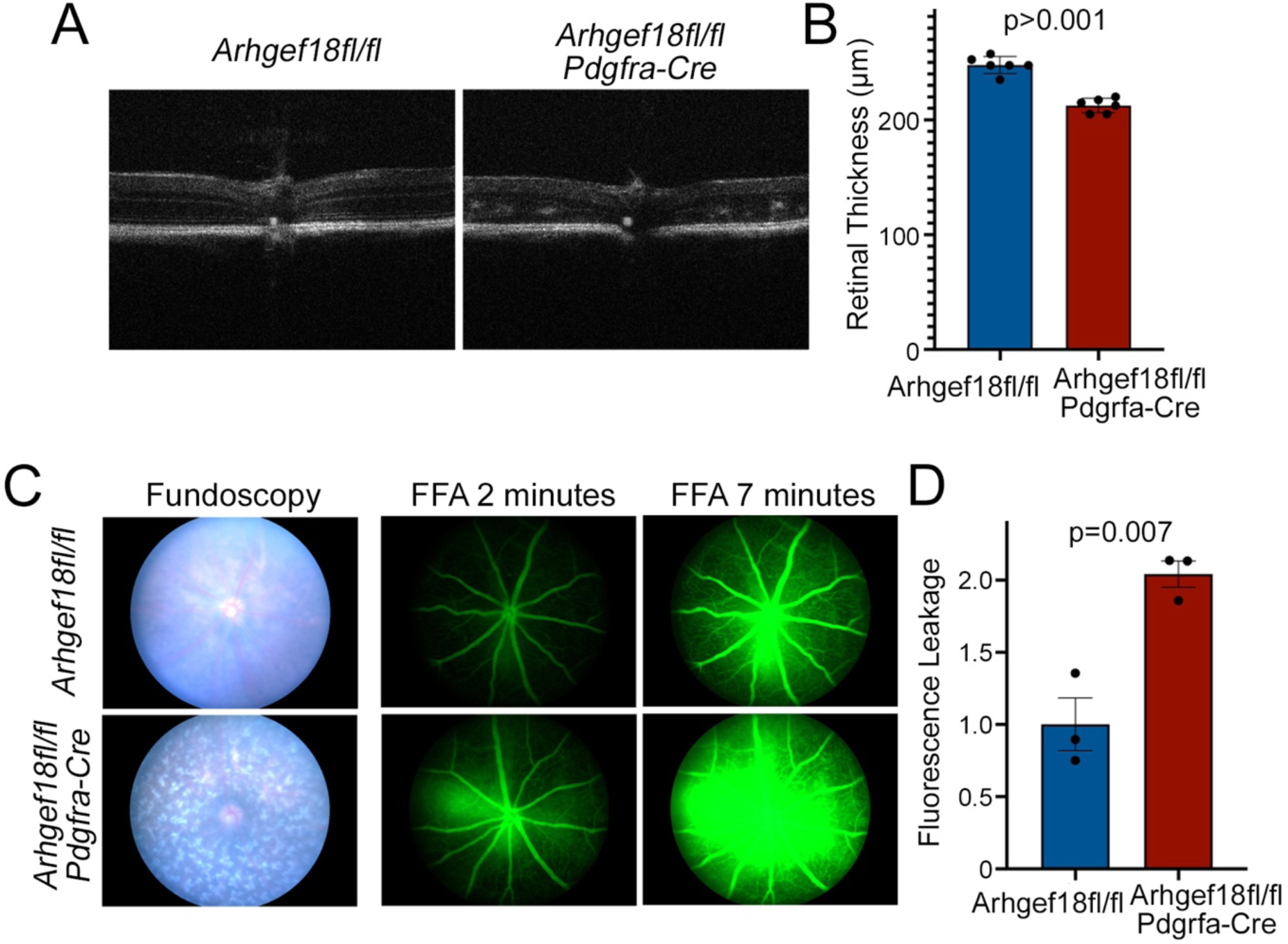
*Arhgef18fl/fl Pdgfra-Cre* mice have reduced retinal thickness and increased retinal vascular leakage. (A) SD-OCT scans from *Arhgef18fl/fl* and *Arhgef18fl/fl Pdgfra-Cre* mice at P21. (B) Quantification of retinal thickness from SD-OCT scans. *Arhgef18fl/fl Pdgfra-Cre* mice showed ∼15% reduction of retinal thickness compared to controls (n=6, shown are means ±1SD; p-values, two-sided t-tests). (C) Fundoscopy and fluorescent angiography (FFA) of *Arhgef18fl/fl* and *Arhgef18fl/fl Pdgfra-Cre* mice at P21. The fundus of *Arhgef18fl/fl Pdgfra-Cre* mice showed many white patchy areas which were not observed in *Arhgef18fl/fl* mice. (D) Quantification of fluorescence leakage. The changes of fluorescence leakage were measured by measuring the increase from 2 to 7 minutes (n=3, shown are means ±1SD; p-values, two-sided t-tests).

The fundus images of *Arhgef18fl/fl Pdgfra*-*Cre* knockout mice displayed many white spots and patchy areas, indicating structural changes in the retina (Fig.2C). FFA revealed that the knockout led to increased blood-retinal barrier leakage (Fig.2C,D). Therefore, Arhgef18/p114RhoGEF expression in MGCs is essential for normal retinal structure and visual function, and its loss led to early onset progressive retinal degeneration including increased vascular leakage.

Arhgef18/p114RhoGEF expression was also detected in the retinal vasculature, colocalising with the junctional markers ZO-1 and p120 catenin (Fig.1A). However, knockout of *Arhgef18* in endothelial cells using a *Tie2-Cre* transgene did not cause any apparent effects on retinal structure, retinal vascular permeability, or ERGs (Fig.S3). This is in line with our previous observation that *Tie2-Cre*-driven endothelial *Arhgef18* knockout animals do not display apparent embryonic defects^6^.

### *Arhgef18fl/fl Pdgfra-Cre* mice develop retinal rosettes

We next performed Haematoxylin and Eosin (H&E) staining of retinal sections from P8 to P60 to identify structural changes induced by *Arhgef18* knockout (Fig.3A). The *Arhgef18fl/fl Pdgfra-Cre* retinas showed a gradual increase in the number of rosettes with age. At P8, small aberrant cell clusters appeared interspersed with photoreceptor segments; however, the retinal layers still appeared intact and regular (Fig.3A). At P10, the aberrant cell clusters progressed into developing rosettes that peaked at P15, and the OLM, the linear separation between the ONL and inner segments (IS) was disrupted (Fig.3A,B). Between P15 and P21, mature rosettes formed with the photoreceptor nuclei disrupting the OPL and the INL (Fig.3A,B). At P21, rosettes became compacted, and the photoreceptor segments were disrupted. From this point on, the RPE layer was detached in all H&E images across the retina and the number of photoreceptor nuclei decreased. At P60, the reduction of retinal thickness was striking (Fig.3A). Thus, *Arhgef18* knockout in MGCs led to the progressive formation of retinal rosettes that started in the central retina at P10, close to the optic nerve, and then extended across the entire retina.

**Figure 3:**
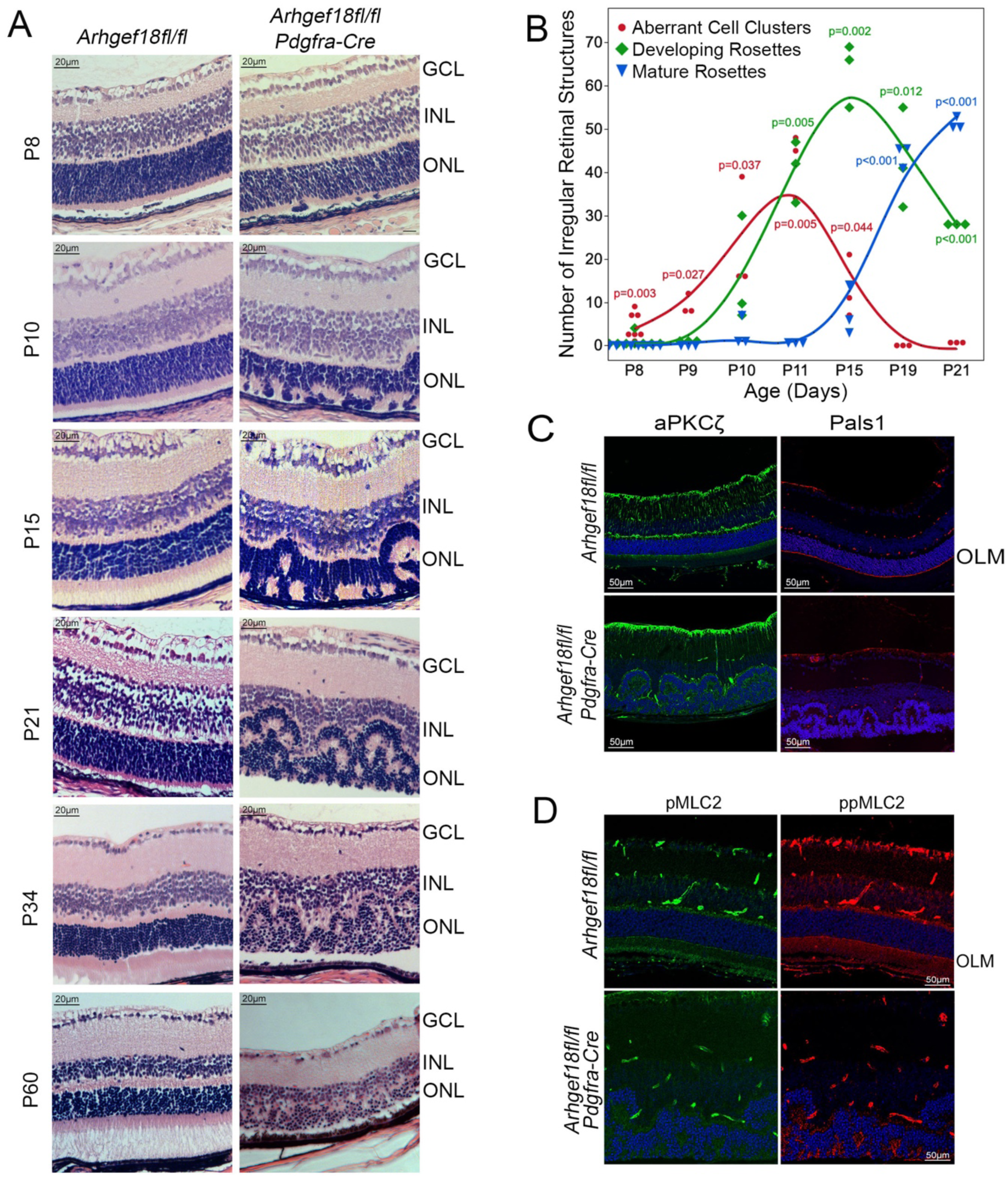
Rosette formation in *Arhgef18fl/fl Pdgfra-Cre* mice. (A) H&E stained images from central retinal sections of *Arhgef18fl/fl* and *Arhgef18fl/fl Pdgfra-Cre* mice *at* indicated ages. ONL (outer nuclear layer); INL (inner nuclear layer); GCL (ganglion cell layer). (B) Quantification of rosettes formation in retinas of *Arhgef18fl/fl and Arhgef18fl/fl Pdgfra-Cre* mice between P8 and P21. Rosettes were counted in 3 sections per eye from 6 mice for P8 or 3 mice for P9 to P21. Note, control *Arhgef18fl/fl* mice did not form aberrant clusters or rosettes (red, aberrant cell clusters; green, developing rosettes; blue line, mature rosettes; shown are datapoints; p-values, one-sided t-tests with a test mean of 0). (C) Expression of OLM-associated polarity markers aPKCζ and Pals1 in retinas of *Arhgef18fl/fl and Arhgef18fl/fl Pdgfra-Cre* mice at P21.(D) Activation of non-muscle myosin-II at the OLM is disrupted in *Arhgef18fl/fl Pdgfra-Cre* mice. Shown is staining for single (pMLC2) and double (ppMCL2) phosphorylated myosin regulatory light chain (MLC2). In panels C and D, nuclei are shown in blue.

The OLM is a large adherens junction between MGCs and photoreceptors at the border between the ONL and inner segments^7^. By P21, the OLM was completely disrupted in *Arhgef18fl/fl Pdgfra-Cre* knockout retinas (Fig.1A). Arhgef18 interacts with the apical polarity signalling machinery, which associates with the OLM in the retina^14,15^. Key components of apical polarity complexes, such as aPKCζ and Pals1, were lost from the OLM of *Arhgef18fl/fl Pdgfra-Cre* mice retinas (Fig.3C). Arhgef18/p114RhoGEF also associates with myosin-IIA and ROCKII, and is a regulator of the actomyosin cytoskeleton^2^. RhoA-stimulated myosin light chain 2 (MLC2) phosphorylation and, thereby, myosin-II activation, is important for stabilization of cell-cell junctions^2–4^. MLC2 can either be mono-(pMLC2) or di-phosphorylated (ppMLC2) in response to RhoA-stimulated ROCK activation; both forms were detected along the OLM in control retinas but lost in *Arhgef18fl/fl Pdgfra-Cre* retinas (Fig.3D). Thus, *Arhgef18* knockout in MGCs led to disruption of components of the apical polarity complexes and myosin activation at the OLM.

At the onset of rosette formation at P10, the junctional markers ZO-1, p120 catenin, and Pals1, were still present at the OLM; however, occasional breaks already started to appear (Fig.4). By P15, when the mature rosettes were fully formed, the OLM was disintegrating in *Arhgef18fl/fl Pdgfra-Cre* retinas. By P21, there remained little detectable staining of the junctional markers ZO-1 and p120 catenin, and the polarity protein Pals1. Thus, Arhgef18/p114RhoGEF in MGCs is essential for stabilization of the OLM and, in its absence, disruption of the OLM correlates with rosette formation and retinal degeneration.

**Figure 4:**
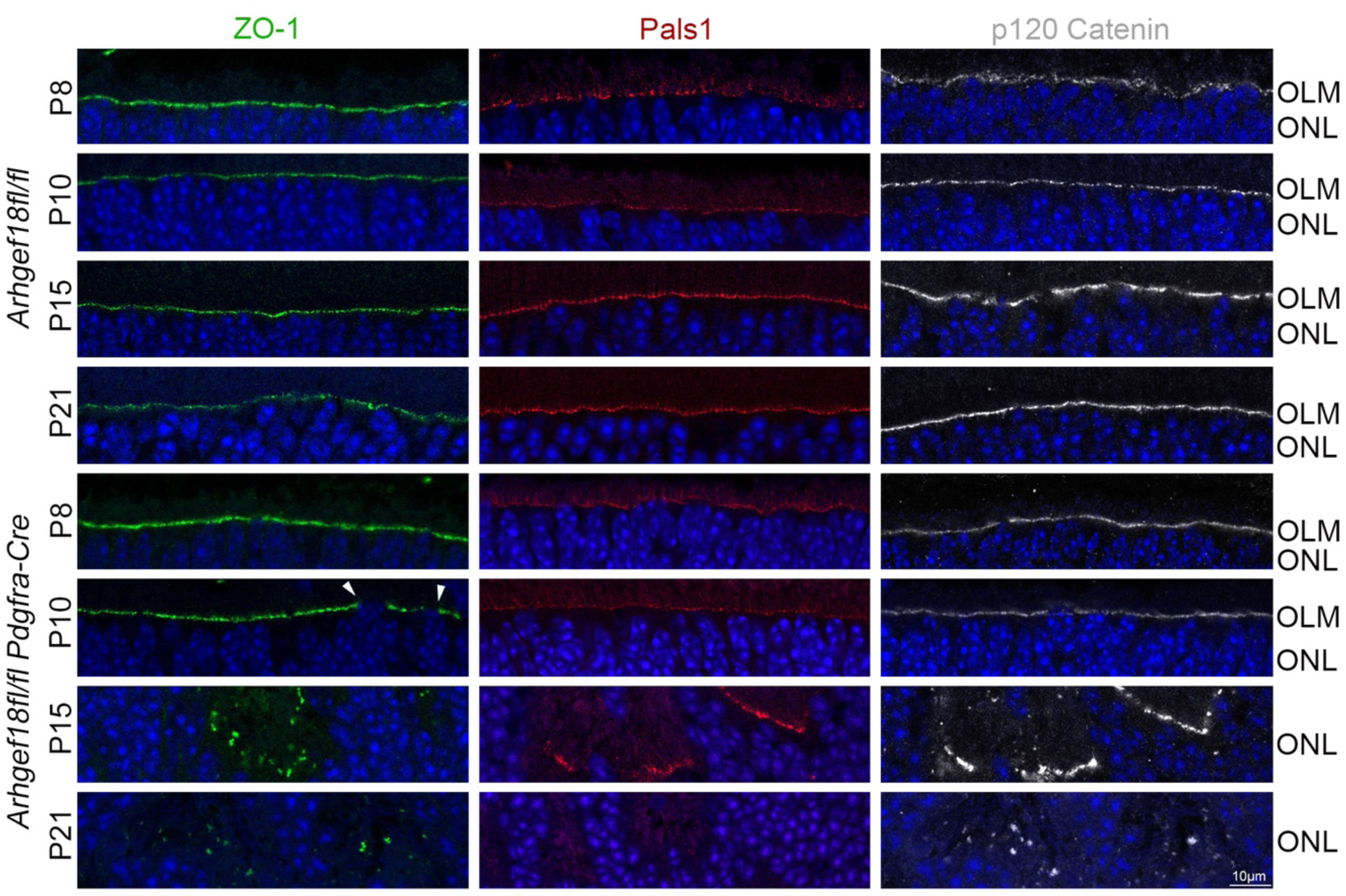
Loss of Arhgef18/p114RhoGEF in MGCs leads to progressive OLM disruption. OLM staining for the junctional proteins ZO-1 and p120 Catenin, and the polarity marker Pals1. Samples were derived from *Arhgef18fl/fl* and *Arhgef18fl/fl Pdgfra-Cre* from P8 to P21. At least three different mice were analysed for the three labels. The positions of the OLM (outer limiting membrane) and ONL (outer nuclear layer) are indicated. Nuclei are shown in blue.

### Loss of Arhgef18/p114RhoGEF in MGC causes photoreceptor cell death

To understand the cell types affected in the retinal degeneration of *Arhgef18fl/fl Pdgfra-Cre* mice, we used retinal cell markers. Bipolar (Chx10) and photoreceptor (Arrestin) cells were disordered, and the latter staining indicated redistribution of segments throughout the rosettes in the ONL of *Arhgef18fl/fl Pdgfra-Cre retinas* at P21 (Fig.5A). The disruption was observed along the entire retina, and it was consistent with other cell type markers such as Sox9, GFAP, vimentin, GS, Recoverin and Rhodopsin (Fig.5B-C, Fig.S4). By P60, little Arrestin staining remained, and the ONL, INL, and OPL were reduced and disrupted (Fig.5A). Quantification of photoreceptor nuclei revealed that control mice maintained ∼1000 nuclei per field of view across timepoints, but *Arhgef18fl/fl Pdgfra-Cre* retinas displayed a significant loss of photoreceptors (∼70%) by P60 (Fig.5A). The nuclei of bipolar cells (Chx10) at the INL were disordered, but the total number was not changed in knockout retinas (Fig.5A). The number of MGCs (Sox9) per field of view was not altered at P21 and P60; however, the cells were dispersed, reflecting the disordered INL in *Arhgef18fl/fl Pdgfra-Cre* retinas (Fig.5B). MGC gliosis, visualised with glial-fibrillary acidic protein (GFAP) increased at P21 and P60 in *Arhgef18fl/fl Pdgfra-Cre* retinas (Fig. 5C). GFAP- and vimentin-positive fibres in *Arhgef18fl/fl Pdgfra-Cre* retinas appeared no longer in a radial patten stretching from the inner towards the outer retina but, by P60, their orientation was strongly disorganised. This was further supported by the staining for glutamine synthase (GS), a constitutive MGC marker, that highlighted the morphological disruption of MGCs in the central and peripheral parts of the retina (Fig.S4). In *Pdgfra-Cre* only retinas vimentin induction or retinal abnormalities was not observed (Fig.S1A). Thus, the retinal degeneration of *Arhgef18* knockout in MGCs affects the adherens junction formed by MGC and photoreceptors and induced gliosis, indicating retinal inflammation, and MGC stress and activation.

**Figure 5:**
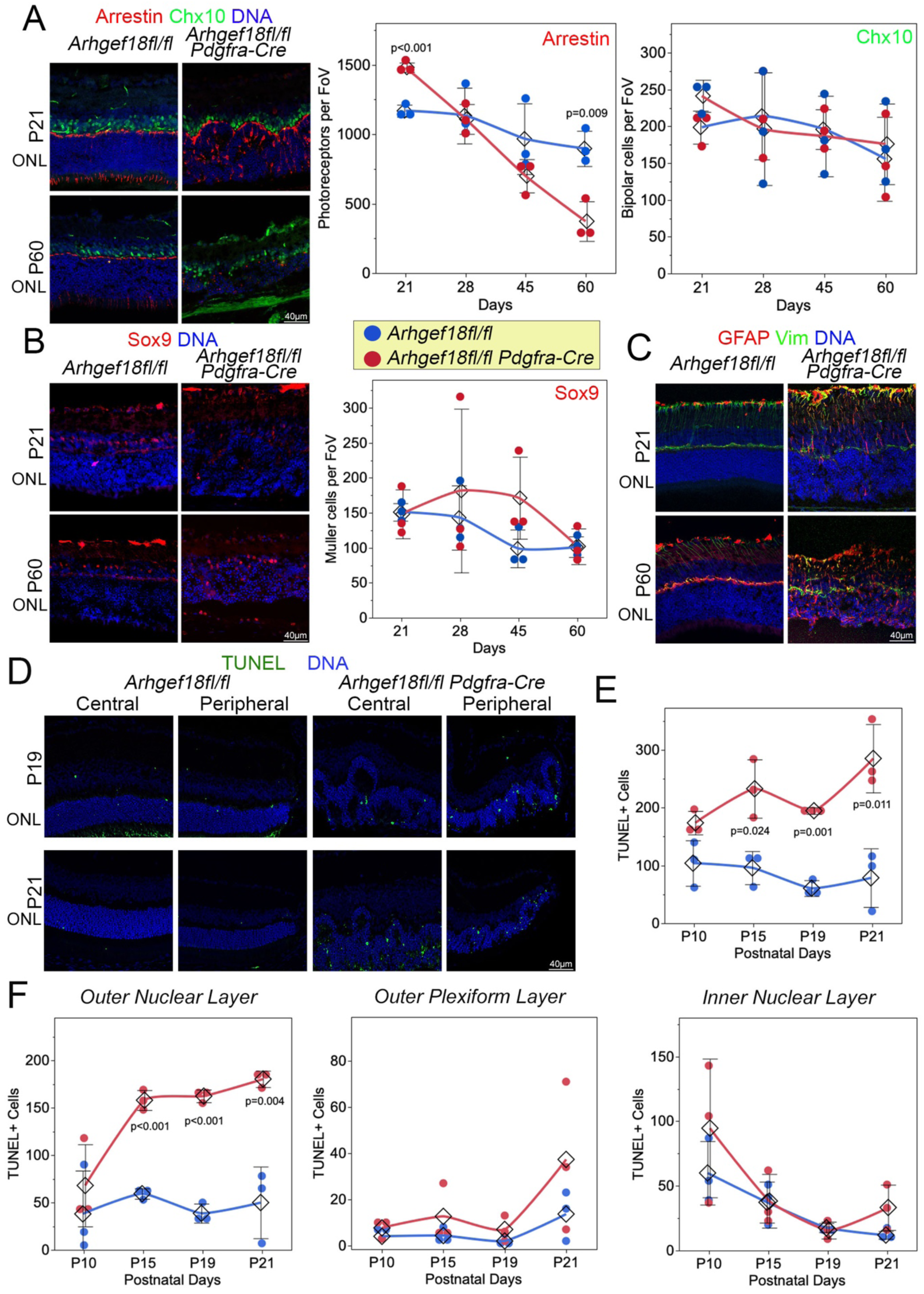
*Arhgef18fl/fl Pdgfra-Cre* mice progressively lose photoreceptors. *(A)* Arrestin, Chx10 and DNA staining of *Arhgef18fl/fl* and *Arhgef18fl/fl Pdgfra-Cre* retinal sections at P21 and P60. Graphs show quantifications of the number of photoreceptors based on Arrestin and Chx10 staining, respectively. Shown are datapoints (n=3), means ±1SD, and p-values from two-sided t-tests. (B) Sox9 and DNA staining of *Arhgef18fl/fl* and *Arhgef18fl/fl Pdgfra-Cre* retinal sections of mice at indicated ages. The quantification shows the number of Sox9-positive MGCs. Shown are datapoints (n=3), means ±1SD, and p-values from two-sided t-tests. (C) GFAP and Vimentin immunofluorescence of *Arhgef18fl/fl* and *Arhgef18fl/fl Pdgfra-Cre* retinas sections from P21 and P60. Nuclei are shown in blue. (D-F) TUNEL assay for the detection of apoptosis in the central and peripheral retina on sections from *Arhgef18fl/fl* and *Arhgef18fl/fl Pdgfra-Cre* mice (green, apoptotic cell; blue, nuclei). Panels E and F show quantifications of TUNEL-positive cells at P10, P15, P19 and P21 either of the entire retina (E) or in specific retinal layers. Shown are datapoints (n=3), means ±1SD, and p-values from two-sided t-tests.

The decreasing retinal thickness and reducing number of photoreceptors is likely to involve cell death. To determine when cell death started, we performed TUNEL assays to identify apoptotic cells (Fig. 5D). Quantification of TUNEL positive cells between P10 and P21 showed increased apoptosis in *Arhgef18fl/fl Pdgfra-Cre* knockout mice across the retina (Fig.5E). The increased in TUNEL positive staining were predominantly associated with the ONL (Fig.5F), indicating that already early apoptotic cells were mostly photoreceptors. Thus, deficiency of Arhgef18/p114RhoGEF in MGCs led to induction of photoreceptor apoptosis already at early stages of retinal degeneration, correlating with disruption of the OLM.

### ARHGEF18/p114RhoGEF downregulation in Müller cells leads to activation of NF-κB and β-catenin/TCF signalling, and mitochondrial dysfunction

ARHGEF18/p114RhoGEF regulates mechanical stress on cell junctions by activating the junctional actomyosin cytoskeleton in response to mechanical tension^2–4,6,16^. In control retinas, ppMLC2 decorated the OLM, co-localising with F-actin (Fig.6A). The ppMLC staining was barely detectable at P10 but then progressively increased until P21, correlating with the increasing mechanical stress on the OLM during postnatal growth of the eye^17^. In *Arhgef18fl/fl Pdgfra-Cre* retinas, the ppMLC2 staining in the area of the OLM was not detectable (Fig.6A-B). Hence, the increase in myosin activation at the OLM of control retinas, occurred at times when the knockout retinas started to degenerate and the OLM to disintegrate, suggesting that Arhgef18/p114RhoGEF-stimualted myosin activation supports retinal integrity.

**Figure 6:**
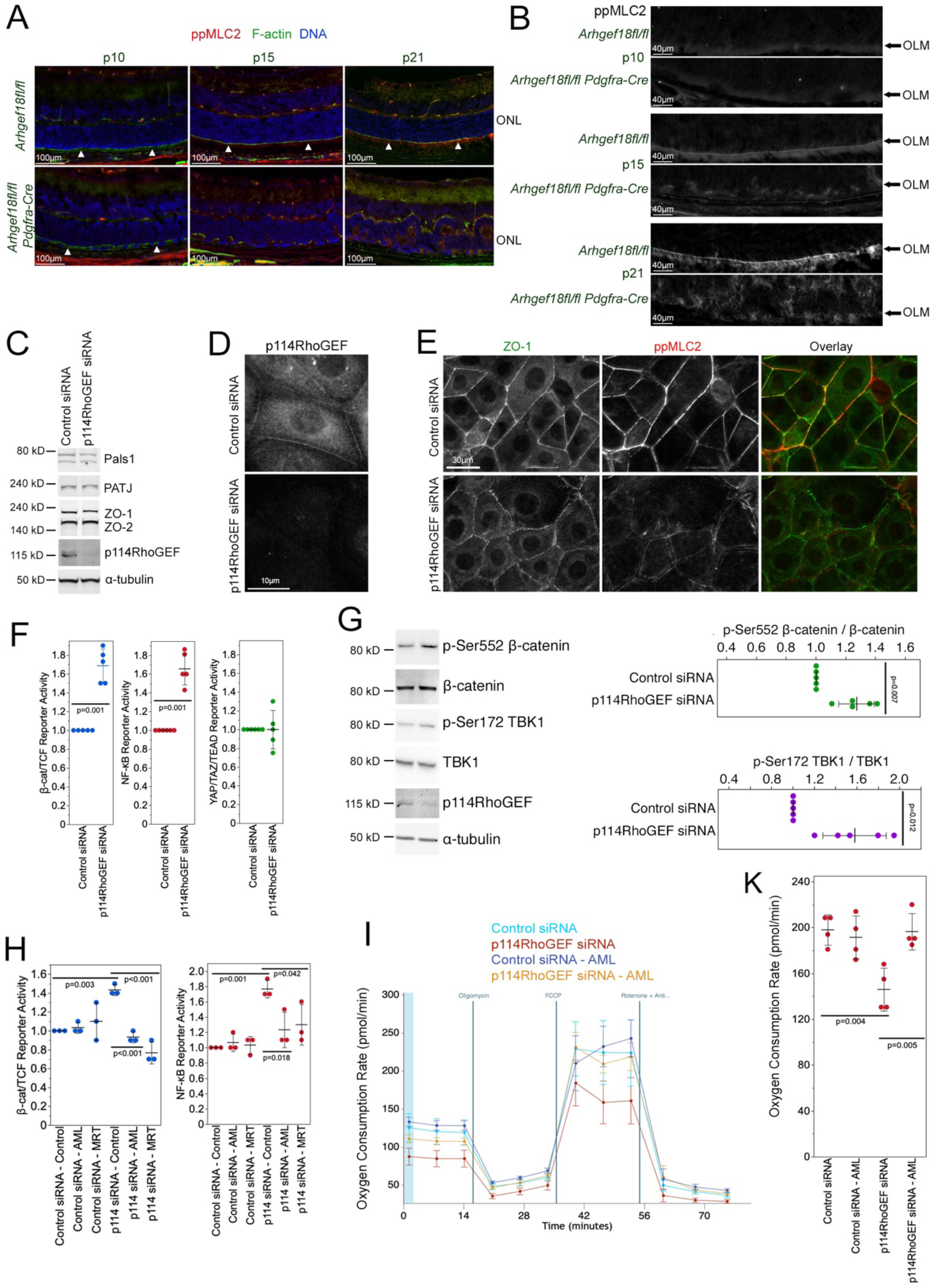
ARHGEF18/p114RhoGEF regulates TBK1-activation and mitochondrial respiration. (A,B) ppMLC, F-actin and DNA staining in retinal sections from *Arhgef18fl/fl* and *Arhgef18fl/fl Pdgfra-Cre* mice at P10, P15 and P21. Panel B shows enlarged parts from panel A images of the ppMLC2 staining. Note, the ppMLC2 staining at the OLM increases with age. (C) Immunoblots of polarity proteins (Pals1, PATJ) and junctional markers (ZO-1, ZO-2) using cell extracts of control and p114RhoGEF siRNA transfected cultured MIO-M1A cells. p114RhoGEF and α-tubulin were blotted to monitor the depletion and total protein loading, respectively. (D-E) Immunofluorescence of cells transfected with siRNAs as in panel C using antibodies for p114RhoGEF (D), or ZO-1 and ppMLC2. (F) β-catenin/TCF, NF-κB, and YAP/TAZ/TEAD reporter gene assays from MIO-M1A cells without or with p114RhoGEF depletion by RNA interference (n=5; shown are datapoints, means ± 1SD; p-values, two-sided t-tests). (G) Immunoblots monitoring phosphorylation of β-catenin at Ser552 and TBK1 at Ser172 in response to p114RhoGEF depletion in MIO-M1A cells. The quantifications show relative increases in phosphorylation normalised to control siRNA transfected cells (n=5; shown are datapoints, means ± 1SD; p-values, two-sided t-tests). (H) β-catenin/TCF and NF-κB reporter gene assays without or with p114RhoGEF depletion in the absence or presence of either one of the two TBK1 inhibitors Amlexanox (AML, 10μM) or *MRT68601 (MRT,* 2μM) (n=3; shown are datapoints, means ± 1SD; p-values, Tukey-Kramer HSD)*. (I-K) Oxygen consumption curves measured with an Agilent Seahorse analyzer using a mitochondrial stress test protocol.* MIO-M1A cells were transfected with siRNAs as indicated and then incubated with the TBK1 inhibitor Amlexanox (AML). The quantification in panel K shows *oxygen consumption rates at maximal mitochondrial respiratory capacity* (n=4; shown are datapoints, means ± 1SD; p-values, Tukey-Kramer HSD).

We next investigated the underlying mechanisms in a cell culture model, using a subclone of MIO-M1A cells that forms cell-cell junctions (Fig.6)^18^. Immunoblotting revealed ARHGEF18/p114RhoGEF depletion by RNA interference and no significant changes in the expression of OLM proteins (Fig.6C, Fig.S5A). ARHGEF18/p114RhoGEF depletion was also detected by immunofluorescence and induced a reduced and zipper-like staining of junctional proteins (ZO-1, N-cadherin, p120 catenin) and a strong reduction in junctional ppMLC2 staining (Fig.6D,E; Fig.S5B). Hence, ARHGEF18 is required for junctional actomyosin activation in MGCs. ARHGEF18/p114RhoGEF also reduced the junctional localisation of the apical polarity proteins Crb2, PKCζ, and Pals1 (Fig.S5C,D). The MGC markers glutamine synthase (GS) and SOX9 were not affected by ARHGEF18/p114RhoGEF, mirroring the *in vivo* data that did not indicate a reduction in MGCs in *Arhgef18fl/fl Pdgfra-Cre* retinas (Fig.5B; Fig.S5D). Hence, loss of ARHGEF18/p114RhoGEF disrupts the normal localisation of junctional and polarity proteins *in vitro* and *in vivo*. Therefore, we used the cell model next to investigate possible underlying mechanisms.

We first performed a pathway profiling screen using different promoters responsive to different transcription factors^19^. Figure 6F shows that depletion of ARHGEF18/p114RhoGEF in MIO-M1A cells stimulated β-catenin/TCF- and NF-κB-dependent promoters. Increased β-catenin/TCF activity was paralleled by increased β-catenin phosphorylation at Ser552 (Fig.6G) and increased cytosolic and nuclear β-catenin staining (Fig.S5D). Ser552 phosphorylation stimulates transcriptional activity and can be phosphorylated by TBK1, a kinase that can also activate inflammatory signalling^20,21^. Indeed, Ser172 phosphorylation of TBK1 was also increased, indicating activation (Fig.6G). Amlexanox and MRT68601 are TBK1 inhibitors used in clinical applications and preclinical studies, respectively^22,23^. Thus, we used the two TBK1 inhibitors to assess if they can attenuate the effects of ARHGEF18/p114RhoGEF depletion in reporter gene assays. Indeed, both inhibitors prevented the induction of β-catenin/TCF- and NF-κB-dependent promoters (Fig.6H). Thus, TBK1 inhibition blocks transcriptional signalling pathways activated by ARHGEF18/p114RhoGEF depletion

In response to injury or disease, Müller cells can become reactive and undergo changes in their metabolic pathways^24^. TBK1 can impact various metabolic pathways which, in turn, regulates mitochondrial activity^25^. Hence, we performed mitochondrial stress tests using a Seahorse analyzer. Indeed, depletion of ARHGEF18/p114RhoGEF decreased maximal and basal respiration, indicating decreased mitochondrial capacity and decreased steady state activity, respectively (Fig.6I,K; Fig.S5E). Both maximal and basal respiration were rescued by Amlexanox. Hence, inhibition of TBK1 rescues mitochondrial activity in ARHGEF18/p114RhoGEF-deficient cells.

Mitochondrial respiration can be stimulated by increasing NAD+ levels with nicotinamide^26^. In agreement, nicotinamide rescued the ARHGEF18/p114RhoGEF-depletion induced decrease in maximal respiration (Fig.7A; Fig.S5F,G). Nicotinamide also attenuated the increase in TBK1 phosphorylation, NF-κB and β-catenin/TCF activation in reporter gene assays, and, as TBK1 inhibitors too, the increased Ser552 phosphorylation of β-catenin, suggesting that stimulating mitochondria can feedback to inhibit TBK1 signalling (Fig.7B-E). Similarly, the junctional disruption induced by ARHGEF18/p114RhoGEF depletion was rescued by the TBK1 inhibitors and nicotinamide (Fig.7F). Hence, TBK1 functions as a mitochondrial regulator in MGCs that is controlled by feedback inhibition. ARHGEF18/p114RhoGEF depletion promotes TBK1 activation and, thereby, induces loss of the equilibrium between TBK1 and mitochondrial activity.

**Figure 7:**
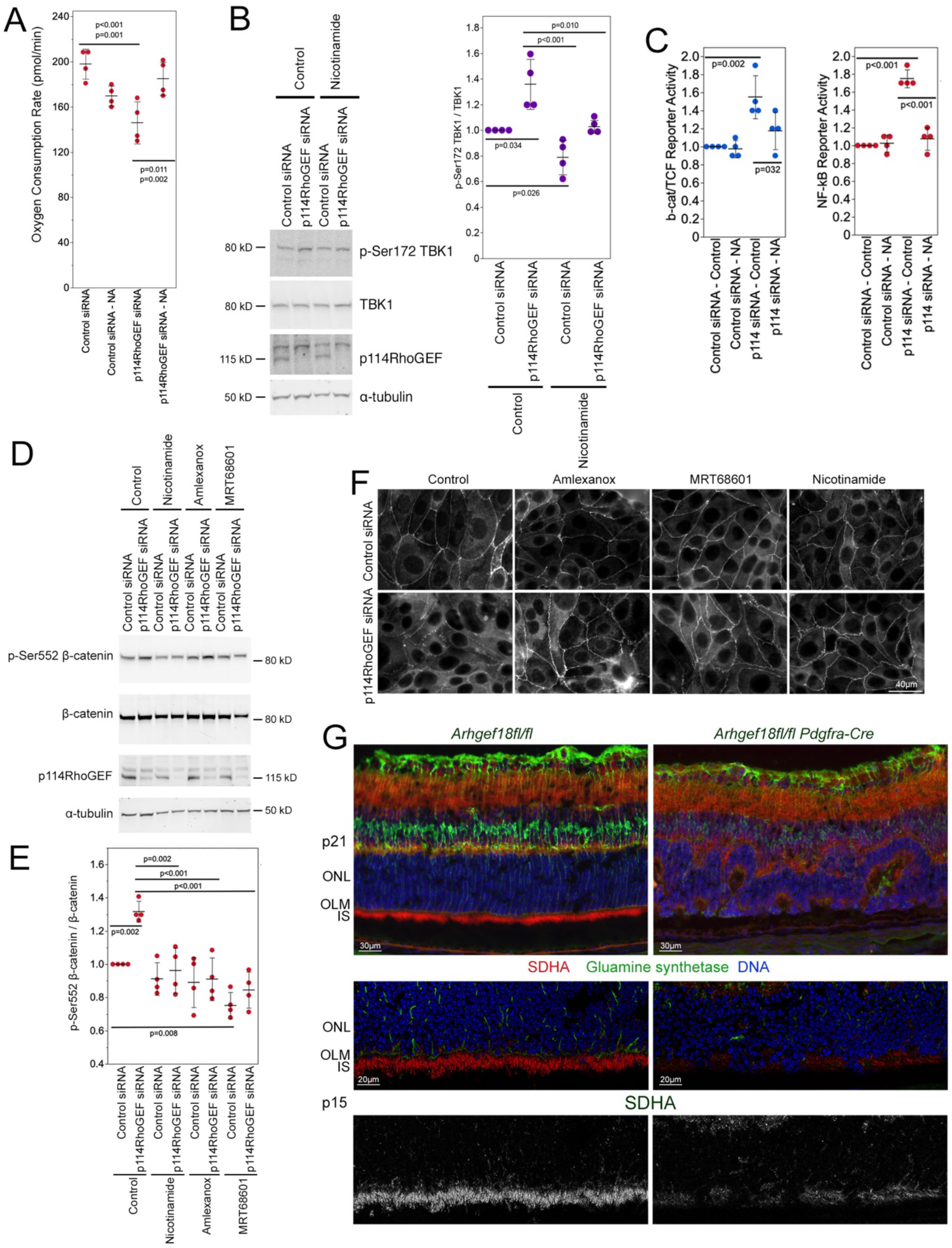
Nicotinamide attenuates ARHGEF18/p114RhoGEF depletion-induced signalling responses. (A) *Oxygen consumption rates were measured during mitochondrial stress tests as in* figure 6 *but cells were incubated without or with nicotinamide (*10μM*). The values without nicotinamide are the same as the controls in* figure 6K *as the experiments were performed in parallel* (n=4; shown are datapoints, means ± 1SD; p-values, Tukey-Kramer HSD). (B) Nicotinamide reduces phosphorylation of TBK1 at Ser172. Immunblots were performed after incubating siRNA transfected cells without or with nicotinamide. The quantification shows datapoints, means ± 1SD; p-values, Tukey-Kramer HSD. (C) β-catenin/TCF and NF-κB reporter gene assays without or with p114RhoGEF depletion in the absence or presence of nicotinamide (n=4; shown are datapoints, means ± 1SD; p-values, Tukey-Kramer HSD). *(D,E)* Immunoblots monitoring β-catenin phosphorylation at Ser552 in cells transfected with siRNAs and incubated without or with TBK1 inhibitors or nicotinamide as indicated. The quantification in panel E shows datapoints, means ± 1SD, and p-values from Tukey-Kramer HSD (n=4). (F) TBK1 inhibitors and nicotinamide rescue junction formation in ARHGEF18/p114RhoGEF-depleted cells. Cells were transfected with siRNAs and then incubated with inhibitors or nicotinamide as indicated. The cells were then fixed and stained for the junctional marker ZO-1 by immunofluorescence. (G) Retinal sections from *Arhgef18fl/fl* and *Arhgef18fl/fl Pdgfra-Cre* mice at p21 and p15 were stained for the mitochondrial marker SDHA and the MGC marker glutamine synthetase by immunofluorescence. Note, control mice display a ribbon of mitochondrial staining in photoreceptor inner segments (IS) adjacent to the OLM that disappears in knockout mice.

We next asked whether mitochondria are affected in *Arhgef18fl/fl Pdgfra-Cre* mice. Specific immunolocalization of mitochondria in MGC *in vivo* is not feasible. However, TBK1 activation has been linked to junction disruption^27^, and photoreceptors form cell-cell junctions with MGC at the OLM. Photoreceptors are packed with mitochondria in their inner segments as illustrated by staining for SDHA (Fig.7G). Mitochondrial localisation is critical for photoreceptor homeostasis and function^28^. Strikingly, with the onset of OLM disruption in *Arhgef18fl/fl Pdgfra-Cre* mice, this retinal ribbon of photoreceptor mitochondria disappeared, suggesting that OLM integrity is tightly linked to the distribution of mitochondria in photoreceptors.

## Discussion

Our data show that ARHGEF18/p114RhoGEF is an essential component of the cell-cell adhesion junction of the OLM, formed by MGCs and photoreceptors. ARHGEF18/p114RhoGEF was not required for OLM formation but for its maintenance and stability. Its deletion in MGCs led to a severe retinal degeneration with vison loss, resembling the phenotypes of patients with biallelic mutations in ARHGEF18 (e.g., retinitis pigmentosa) but starting earlier. Unlike the knockout mouse model, the identified mutations in patients are only partially inactivated (i.e., reduced catalytic activity or active but mislocalised)^1^; hence, one would expect gene knockout in mice to be more severe or to promote disease in younger animals. The OLM of the Arhgef18-knockout retinas started to form rosettes by P10, correlating with the onset of OLM destabilisation, and, by P21, reached a severe retinal degeneration with vascular leakage. In cultured human MGCs, ARHGEF18/p114RhoGEF depletion stimulated a TBK1-dependent signalling mechanism that led to reduced mitochondrial respiration. Both inhibition of TBK1 or direct stimulation of mitochondria attenuated downstream signalling mechanisms and inhibition of mitochondrial function.

Arhgef18 localises at the OLM, and MGC knockout impairs visual function. The OLM is a continuous heterotypic cell-cell adherens junctions between Müller cells and photoreceptors^29,30^. Here, we discovered that ARHGEF18/p114RhoGEF is an essential component of the OLM. Active ARHGEF18/p114RhoGEF stimulates the actomyosin cytoskeleton, resulting in stabilisation of intercellular junctions. In the OLM, ARHGEF18/p114RhoGEF was not required in MGCs for OLM formation but for OLM stabilisation after MGC integration into the retina. The onset of OLM destabilisation in knockout mice was when phosphorylated MLC2 started to appear in wild-type mice, which was lost in *Arhgef18fl/fl Pdgfra-Cre* mice, indicating that the increase in active myosin was ARHGEF18/p114RhoGEF-dependent. ARHGEF18/p114RhoGEF regulates mechanical stress on cell junctions, stabilising the junction by activating the actomyosin at the cell-cell contact sites in response to mechanical tension^2–4,6,16^. The increase in MLC2 phosphorylation at the OLM occurred at a time when the mechanical stress on the OLM increased due to the postnatal growth of the eye^17^. Hence, our data indicate that the function of ARHGEF18/p114RhoGEF is to support the role of MGCs in maintaining the mechanical stability of the retina^31^.

*Arhgef18 Pdgfra-Cre* retinas at P21 exhibit nummular pigmentation similar to what it is observed in retinitis pigmentosa (RP) of patients carrying biallelic mutations in *ARHGEF18*^1^. The disruption of the OLM in *Arhgef18fl/fl Pdgfra-Cre* retinas started at P8, after integration of MGCs into the retina, and the OLM disruption led to loss of retinal thickness due to photoreceptor apoptosis, retinal degeneration, and visual impairment by P60. Patients with biallelic mutations in ARHGEF18 have adult-onset retinal degeneration, characterized by retinal changes resembling those caused by *CRB1* mutations, including intraretinal cysts and RP. *Crb1* knockout in mice leads to late onset retinal degeneration that worsens upon white light exposure^10, 32^. Loss of *Crb2* in MGCs in *Crb1* knockout mice accelerates the phenotype and, like *Arhgef18*, leads to retinal degeneration more quickly^7^. However, the induced retinal phenotypes are morphologically different, and OLM defects are observed at E17.5 prior to MGC integration into the retina in the *Crb1/2* model; hence, direct comparisons are difficult to make. *Arhgef18* expression in Müller cells is essential for maintenance of intercellular adhesion between Müller glia and photoreceptors, retinal structure, and function. However, it is possible that Crb1 and Crb2 play a role in regulating ARHGEF18/p114RhoGEF similar to its Drosophila homologue Cysts that is regulated by apical polarity proteins. Nevertheless, junctional components are important for ARHGEF18/p114RhoGEF recruitment and function in vertebrate cells under tension^3,4,33^. As the MGC *Arhgef18* knockout mice assemble apparently morphologically normal retinas, and the OLM then disassembles when mechanical stress increases, ARHGEF18/p114RhoGEF’s function in MGCs seems to be primarily connected to its role in mechanical stabilisation of cell-cell adhesion rather than establishment of apicobasal polarity.

*Arhgef18fl/fl Pdgfra*-*Cre* mice also displayed increased retinal vascular permeability, which had not been reported in other mouse models of OLM disruption^7,12,34^, suggesting that defects in MGCs due to *Arhget18* depletion play a role in cross-regulation of retinal endothelial cell functions. Knocking out *Arhgef18* with a *Tie2-Cre* transgene, which knocks out floxed genes in endothelial cells, did not lead to retinal vascular permeability or detectable changes in retinal structure or function. Hence, the knockout in MGCs may lead to vascular stress and, thereby, increased vascular permeability. The *Crb1/2* mutant mice mentioned above display signs of neovascularisation^7^; hence, vascular stress may be a common feature of OLM-disruption associated retinal degenerations. Once possibility is that the reduced mitochondrial function we observed in ARHGEF18/p114RhoGEF-depleted cells may lead to signals that induce retinal vascular stress.

The available antibodies for Crb1 and Crb2 did not work in our mouse tissue sections, and only Crb2 antibodies worked for cultured human MGCs. Immunofluorescence indicated only a modest effect on Crb2 localisation upon ARHGEF18/p114RhoGEF-depletion but cytosolic polarity proteins like Pals1 and aPKCζ were more severely affected. Similarly, polarity proteins like *Pals1* were strongly affected in *Arhgef18 Pdgfra-Cre* retinas. *Pals1* knockout in retinal progenitors leads to the disruption of the apical localization of Crb proteins in retinal progenitors, early visual impairments, and disorganization of retinal lamination^35^. Our data indicate that *Arhgef18* is required for maintaining proper localisation of Pals1 at the OLM, indicating that Arhgef18 may function as a modulator of the Pals1/Crumbs complex by supporting the maintenance of cell-cell adhesion and, thereby, maintenance of apical polarity, as the redistribution of Pals1 only occurs with destabilisation of the OLM. Hence, Arhgef18 is not required to establish cell polarity and the OLM but for their maintenance.

ARHGEF18/p114RhoGEF depletion in Müller cells in culture led to activation of NF-κB and β-catenin/TCF signalling. In the retina, NF-κB signalling is activated in Müller cells after neuronal damage and during chronic degeneration, promoting glial reactivity^36^. Similarly, β-catenin/TCF signalling in MGCs plays a crucial role in retinal health and disease by influencing neuronal regeneration, photoreceptor protection, and retinal development^37,38^. Hence, the two pathways may have been activated in MGCs as a general response to counteract retinal damage. In the cell culture model, we also measured activation of TBK1, and its inhibition attenuated activation of NF-κB and β-catenin/TCF signalling. We previously observed TBK1 activation in response to ZO-1 KO in epithelial cells, a manipulation that leads to destabilisation of tight junctions^27^ including p114RhoGEF miss-localisation. Hence, activation of TBK1 may be a common mechanism triggered by dissociation of intercellular junctions.

Our observations link ARHGEF18/p114RhoGEF to TBK1 activation and to the regulation of mitochondrial respiration, which was downregulated in response to ARHGEF18/p114RhoGEF depletion. Metabolic pathways are affected in Müller cells in response to disease^24^. TBK1 can impact energy metabolism and mitochondrial function^25^. Mitochondrial respiration is reduced in ARHGEF18/p114RhoGEF deficient MGCs, which can be rescued by TBK1 inhibition. While it is currently not clear how TBK1 activation promoted downregulation of basal and maximal mitochondrial respiratory activity, NF-κB activation can suppress mitochondrial respiration and oxidative phosphorylation. Since NF-κB can interact with β-catenin signalling and TBK1 inhibition also downregulated the two transcriptional pathways, NF-κB and β-catenin/TCF signalling may function as the effector mechanisms by which ARHGEF18/p114RhoGEF via TBK1 impacts on mitochondria in MGCs^39^. Metabolic defects in a mouse model of retinal degeneration induced by mtDNA mutations serve as a paradigm for mitochondrial retinopathy. They are associated with reduced expression of SDHA and cause retinal cell-specific deficiencies, especially in retinal ganglion cells, rods, and Müller cells^40^. Mitochondria as visualised by SDHA staining are strongly affected in retinas of *Arhgef18fl/fl Pdgfra-Cre* knockout mice. Mitochondria form a ribbon in inner segments of photoreceptors that spreads across the retina. In *Arhgef18fl/fl Pdgfra-Cre* knockout retinas that ribbon was lost with the onset of OLM destabilisation, suggesting that OLM integrity is tightly linked to mitochondrial maintenance in inner segments of photoreceptors. Given the importance of mitochondrial distribution and activity for photoreceptor survival and function, the impact on mitochondria in photoreceptors may play an important role in the retinal degeneration induced by knockout of *Arhgef18* in MGCs.

Nicotinamide, a form of vitamin B3, plays a crucial role in cellular metabolism and has been shown to increase oxidative phosphorylation, and alter mitochondrial morphology and activity in retinal ganglion cells, which results in a neuroprotective effect^41^. Nicotinamide rescued the reduced mitochondrial activity in cultured MGCs. Nicotinamide also downregulated TBK1 signalling, indicating that mitochondrial activity and TBK1 regulation are interconnected by regulatory feedback. Hence, it will be important to test if nicotinamide and/or TBK1 inhibitors can rescue retinal degeneration in *Arhgef18fl/fl Pdgfra-Cre* knockout mice to determine if such treatments may be transferable to clinical use. Both nicotinamide and TBK1 inhibitors have either already been used for clinical applications or are in clinical trials; hence, if successful, such treatment strategies could be translated efficiently.

## Materials and Methods

### Animals

The *Arhgef18* floxed line (Arhgef18^tm1a(KOMP)Mbp^) was purchased from UC Davis KOMP Repository, maintained as homozygotes, and crossed for more than 8 generations in C57BL/6J from Charles River Laboratories UK. We have previously described *Arhgef18fl/fl and Arhgef18fl/fl Tie2-Cre mice*^6^. *Arhgef18* floxed mice (*Arhgef18fl/fl*) were then crossed with mice expressing the *Cre* recombinase downstream of the *Pdgfra* gene promoter (*Pdgfra-Cre*, *C57BL/6-Tg (Pdgfra-Cre)1Clc/J,* Strain: 013148*, The Jackson Laboratory*), a transfene previously used to knockout genes in MGCs^7^. *Arhgef18fl/fl* and *Pdgfra-Cre* mice do not display a phenotype in the retina. *Arhgef18fl/fl* littermates were used as controls in all Cre-indued knockout experiments. Mice were maintained in the Biological Research Unit of the Institute of Ophthalmology, University College London. All experiments conducted were in accordance with the UK Home Office Animals Scientifics Procedures Act 1986 (PPL PP6039705). Animals were maintained on 12 hours light-dark cycle.

### Genotyping

Mouse DNA was obtained from ear samples and genotyped by PCR for the wild-type and the floxed *Arhgef18* alleles using the primers Arhgef18-For1 (5’-GAAGCCAGTTTAAGTGTGCCACTTTGG-3’) and Arhgef18-Rev1 (5’-GTCTGTATGAGTGTGTGTTCATCAC-3’). To genotype *Pdgfra-Cre* mice, we used the primers Cre-F (5’-GCCTGCATTACCGGTCGATGC-3’) and Cre-R (5’-CAGGGTGTTATAAGCAATCCCC-3’). PCR reactions were performed using Megamix Blue master mix (Microzone). The first denaturation step was at 95°C for 3 mins followed by three cycles of 94°C for 15 seconds, 65°C for 30 seconds and 72°C for 40 seconds, and 30 cycles of 94°C for 15 seconds, 55°C for 30 seconds and 72°C for 40 seconds. PCR products were separated on 2% agarose submarine gels.

### Tissue Collection

Enucleated eyes were either fixed in 4% formaldehyde for 24 hours or 4% paraformaldehyde (PFA) in phospho-buffered saline (PBS) for 30 minutes. Formaldehyde-fixed eyes were paraffinized and embedded in paraffin blocks. Formalin-fixed paraffin embedded (FFPE) eyes were sectioned at 4μm thickness. PFA-fixed eyes were cryoprotected in 20% sucrose overnight at 4°C and then embedded in optimal cutting temperature (OCT Embedding matrix, Cellpath, UK) solution and sectioned at 10μm. All eye globes were orientated and embedded in the same position to ensure that sections had the same eccentricity.

### Immunofluorescence Microscopy

For FFPE sections, slides were deparaffinised with Histoclear II (3 × 8 min; National Diagnostics, USA) followed by gradual rehydration rehydration through graded ethanol solutions (100% ethanol: 2 × 6 min; 95%, 90%, 80%, and 70% ethanol: 5 min each) and MilliQ water (5 min). Slides were incubated at 98°C in Universal HIER antigen retrieval reagent (ab208572, Abcam) for 20 minutes, followed by rinsing with PBS and blocking. Primary antibody was incubated in 0.3% Triton X-100 in PBS overnight (see Supplementary Table 1 for primary antibodies used). The sections were washed three times in 0.1% Tween20 in PBS (PBS-Tween) and then incubated with secondary antibodies (diluted 1:700) and Hoechst 33342 for half a day. The slides were then washed three times with PBS-Tween and once with PBS prior to mounting with ProLong Gold (P36930, Thermo Fisher Scientific) and storage at 4°C.

Cryosections were permeabilised with 0.5% Triton-X-100 and 0.1% Saponin in PBS (TSP) and then blocked with 1% Donkey serum in TSP. Primary antibodies were diluted in blocking solution and incubated with the sections overnight. The antibodies used are listed in Supplementary Table 1. The sections were three times washed in five times diluted TSP (diluted with PBS) and then incubated in secondary antibodies solution (TSP, 1:700 diluted secondary antibodies) with Hoechst 33342 for half a day. Following a further three washes in diluted TSP solution and one wash in PBS, the slides were mounted with ProLong Gold. For cell staining, coverslips in 48-well plates were used to culture the cells. The cells were fixed with 3% PFA in PBS for 20 minutes at room temperature and then permeabilised with 0.3% Triton X-100 in PBS-BSA (PBS containing 0.5% BSA and 20mM glycine) for 10 minutes.

The cells were then washed with PBS-BSA and incubated with primary antibodies (Supplementary Table I) overnight. After 3 washes with PBS-BSA, the cells were incubated with secondary antibodies in the same solution. After 4 hours, the cells were washed three times with PBS and mounted on glass slides with ProLong Gold. Images were captured with A Nikon Eclipse Ti-E epifluorescence microscope (using Nikon S Plan Fluor 20× (N.A., 0.45) or 60× oil (N.A., 1.4) objectives and NIS Elements Version 4.60 software) or a Leica SP8 confocal scanning microscope (63× (N.A., 1.40) oil objective and LAS X Version 3.5.7.23225 software).

### Histological Analysis

FFPE sections were used for haematoxylin and eosin staining (H&E) using a Leica ST 5010 Autostainer XL (Leica, Germany). Rosettes were quantified manually using a Leica DM IL inverted microscope. Retinal thickness was measured by imaging the H&E sections either side of the optic nerve (ON). Four images per retina were taken in total and quantified with Image J/Fiji - Version 2.16.0/1.54.

### Terminal deoxynucleotidyl transferase dUTP Nick-End labelling (TUNEL) assay

Cells undergoing apoptosis were detected in FFPE sections by TUNEL assay using the manufacturer provided protocol (G3250; Promega). Images shown in the figure were captured using a Leica SP8 confocal scanning microscope. TUNEL positive marks were quantified manually from images obtained with the Nikon Eclipse Ti-E epifluorescence microscope.

### *In vivo* eye assessments

Mice were injected intraperitoneally with anaesthesia (Ketamine/Dormitor 30µg/5µg per g body weight) and pupils were dilated using topical mydriatic solution (Pharma Stulln, #ATH0713). Eyes were then imaged using spectral-domain optical coherence tomography (SD)-OCT (Spectralis, Heidelberg engineering). Funduscopy and angiography were next performed using a Micron III (Phoenix Micron, Phoenix Technology Group). After funduscopy, mice were subcutaneously injected with 1% Fluorescein (Sigma-Aldrich, #F6377). Fundus fluorescent angiography (FFA) images were then obtained at 2 and 7 minutes post-injection. After imaging, eyes were either collected for histological exanimation or mice were injected intraperitoneally with Antisedan (Vetoquinol) to stimulate recovery.

For electroretinography (ERG), mice were dark-adapted overnight and then anaesthetised with Ketamine/Dormitor as described above. Full-field scotopic ERG was carried out using the Diagnosys system and Espion software (Diagnosys, Lowell, MA) under red-light conditions, as previously described^42^. Briefly, simultaneous bilateral recordings were taken using first scotopic ERG protocols followed by photopic measurements after a 10-minute light adaptation. Flash stimuli (1–10 msec duration) were presented via an LED stimulator (scotopic measurements: 0.005 to 50cds/m^2^; photopic measurements: 0.1 to 50cds/m^2^). ERG responses were collected with the Espion software for analysis. The A-wave amplitude was measured by calculating the difference in amplitude between time 0 (i.e., time of light stimulus) and the minimum. The B-wave amplitude was measuring by calculating the difference between the minimum (the one used for A-wave calculations) and the highest point of the curve. The A-wave implicit time was measured by calculating the time passed between time 0 and the lowest peak point (i.e., the A-wave minimum).

### Cell Culture, siRNA Transfections and Immunoblotting

The MIO-M1A cells were cultured in low glucose Dulbecco′s Modified Eagle′s Medium (DMEM) (Sigma-Aldrich, USA) supplemented with 10% Fetal Bovine Serum, 1% Penicilin-Streptomysin (Sigma-Aldrich, USA), and 10mM Hepes pH 7.4 at 37°C in a 5% CO₂ incubator. The cells were split once a week at a 1 to 10 dilution. Junction-forming cells were isolated from the large islands of cells that formed in monolayers of the original line that has a more mesenchymal appearance. The cells were tested for expression of OLM adhesion proteins and MGC markers (Fig.6 and S5). For siRNA transfections, cells were plated in plates appropriate for the experiments (i.e., 48-well plates with coverslips for immunofluorescence, 12-well plates for the preparation of cell extracts for immunoblotting, 96-well plates for reporter gene and seahorse assays). The cells were then transfected with siRNAs using Lipofectamine™ RNAiMAX (Invitrogen, USA) following the manufacturer’s protocol nontargeting siRNA controls 5′-UGGUUUACAUGUCGACUAA-3′ and 5′-UGGUUUACAUGUUGUGUGA-3′, and human ARHGEF18/p114RhoGEF siRNAs 5′-UCAGGCGCUUGAAAGAUA-3′ and 5′-GGACGCAACUCGGACCAAU-3′^6^. All siRNAs were synthesised with dTdT 3’-overhangs. For immunoblotting, cells were washed once in PBS prior to lysis in SDS-PAGE sample buffer (4x LDS sample buffer, MPSB-250ML, Sigma-Aldrich, USA) and separation of samples on surePAGE™ Bis-Tris 4-12% gradient gels (GenScript Biotech Ltd). Proteins were then transferred to Immobilon-FL PVDF membranes (Merck Life Science) and incubated with primary and secondary antibodies as described in results sections (see Supplementary Table I for antibodies)^27^.

### Reporter Gene Assays

Plasmids containing a promoter with binding sites for specific transcription factors to be measured driving firefly luciferase expression, were co-transfected with a renilla luciferase reporter plasmid with a CMV promoter (Promega Corp, Madison, WI, USA) using TransIT reagent (0.25 μL/well, 96-well plate, and 1 μL/well, 48-well plate, Mirus Bio, Madison, WI, USA). The following day, the transfection mix was replaced with fresh medium and after 2 h the two luciferases were measured sequentially using the dual luciferase assay kit (Promega Corp, Madison, WI, USA) and a BMG FLUOstar OPTIMA microplate reader (Ortenbert, Germany). The TEAD-reporter plasmid was kindly provided by Stefano^43^; the TCF-reporter (TOPFLASH) and a corresponding control plasmid (FOPFLASH) were purchased from Upstate Biotechnology Inc. (Lake Placid, NY, USA)^44^.

### Seahorse Assays

Muller cells were seeded into a Seahorse XFe96/XF Pro Cell Culture Microplate (Agilent, USA) at a density of 0.7*10^4^ cells/well in 50ul of culture medium. The following day the cells were transfected with siRNAs for 24 hours followed by incubating in the fresh culture medium for 48-72 hours. One hour prior to the Seahorse assay, the culture medium was replaced with Seahorse XF Assay Medium, composed of Seahorse XF DMEM Medium (pH 7.4, Agilent, USA) supplemented with 10 mM glucose, 1 mM pyruvate, and 2 mM L-glutamine (Agilent, USA). All experiments measuring energy metabolism were run with a Seahorse XF Pro Analyzer using the software to run and analyse the experiments provided by the manufacturer (Agilent, USA). The mitochondrial stress test was run according to the manufacturer’s instructions using the reagents provided with the respective kits for these assays or components purchased from Merck Life science UK (rotenone, R8875; antimycin A, A8674; FCCP, C2920; oligomycin, 495455). Assay results were normalized by quantifying cell numbers using the CyQUANT assay. Briefly, cells in the Seahorse Cell Culture Microplates were frozen overnight at −80°C. The CyQUANT assay was performed using the CyQUANT™ Cell Proliferation Assay Kit (Invitrogen, USA), following the manufacturer’s guidelines. Fluorescence was measured at an excitation wavelength of 480 nm and an emission wavelength of 520 nm. The CyQUANT data were uploaded into the Seahorse Wave software for normalisation of the Seahorse experiments.

### Statistical Analysis

Images were analysed using Image J/Fiji - Version 2.16.0/1.54. Photoreceptor cell numbers were quantified from Hoechst-stained sections; bipolar and Müller glia cell numbers were based on Chx10 and Sox9 staining, respectively. For photoreceptor cell quantification the outer nuclear layer was selected based on the nuclear staining, and all other retinal layers were removed. This was not needed for Chx10 and Sox9 staining of bipolar and Müller cells, respectively. In all cases, a series of processing filters were used to individualise DNA-stained nuclei, Chx10- and Sox9-positive cells, consisting of Gaussian Blur (sigma=2), Unsharp Mask (radius=4, mask=0.6) and Maximum (radius=2). Then, filtered images were analysed using Stardist2D plugin with the following parameters: Model=Versatile (fluorescent nuclei); Normalize image, Percentile low=2.7; percentile high=99.8; Probability=0.05; Overlap Threshold=0.2; Output Type=Both; ROI position=Stack. Retinal thickness from SD-OCT scans, was quantified using Image J/Fiji, tracing the retinal length at 120 pixels from the optic nerve head, and taking two measurements (right and left of the optic nerve head) per scan to minimise the error. Quantitative data are shown as datapoint and/or means +/− one standard deviation. The n number is provided in the respective figure legends and refers to mice in the *in vivo* studies or independent experiments in the cell-based assays. Statistical analysis was performed with JMP Pro 18. Statistical significance of data pairs was assessed with t-tests and, in experiments requiring multiple comparisons Tukey-Kramer HSD tests following an ANOVA test. Tests were two-sided except in figure 3B, which monitored inductions of structures that only occur in mutant eyes and, hence, compared the numbers observed in mutants to zero.

## Acknowledgements

This project was supported by Fight for Sight UK (PhD fellowship AACF, 5071-5072), Medical Research Council (MRC, MR/W028735/1), Moorfield Eye Charity (GR000029), the BBSRC (BB/X000575/1) and Biomedical Research Centre (BRC) at Moorfields Eye Hospital and the UCL Institute of Ophthalmology.

## Author contributions

AACF, TL, HT, and XF performed investigation, data curation and formal analysis. AACF, TL, HT, and XF also contributed to the original draft of the manuscript. KM and MSB conceived the study and methodology, were responsible for funding acquisition, project administration and superviseon, performed investigation, data curation, and formal analysis, and wrote the manuscript with the support of all authors.

## Competing interests

The authors declare no competing interests.

## Data Availability

The datasets generated and/or analysed during the current study are available from the corresponding author upon reasonable request.

---

**Supplementary Table S1:**
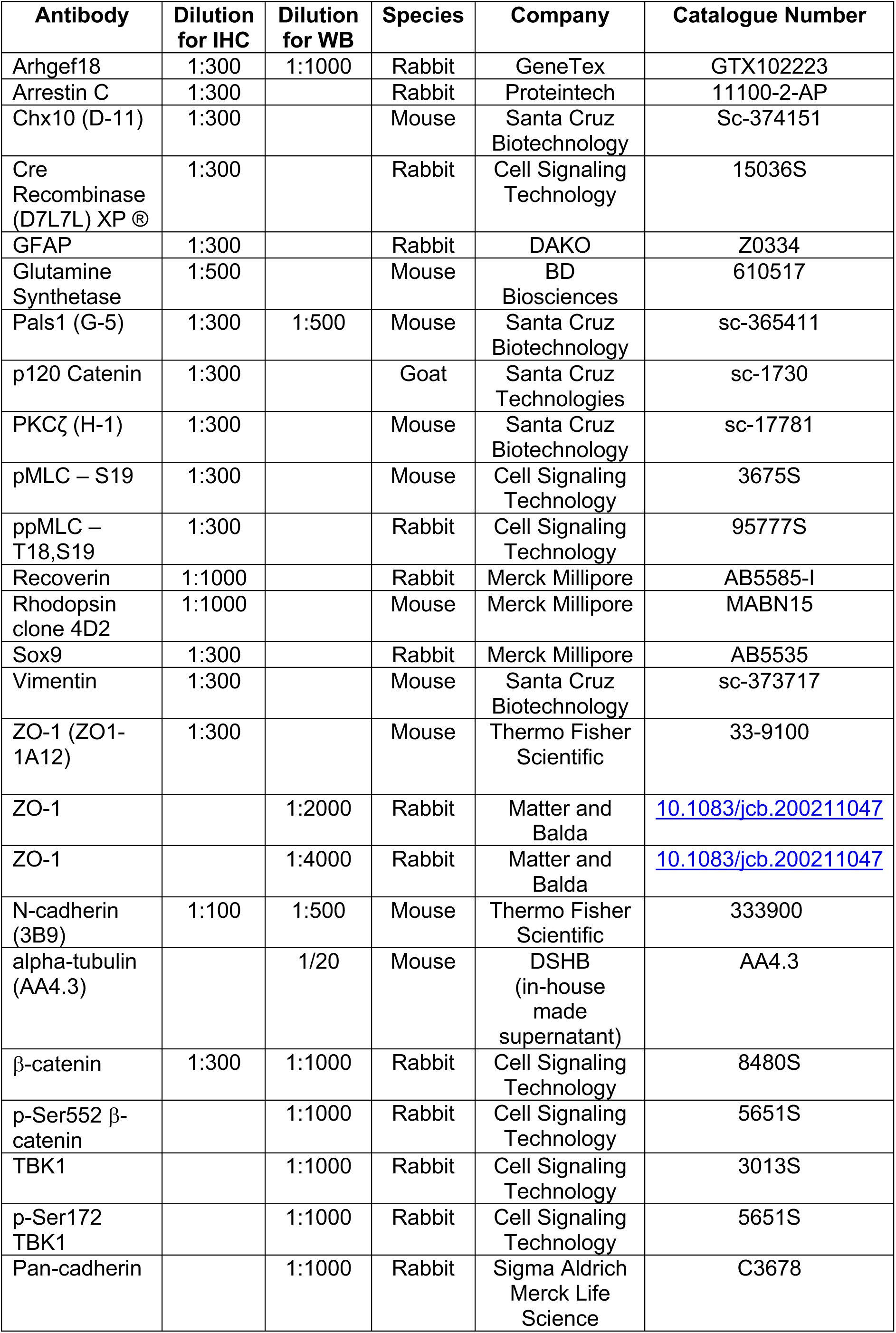
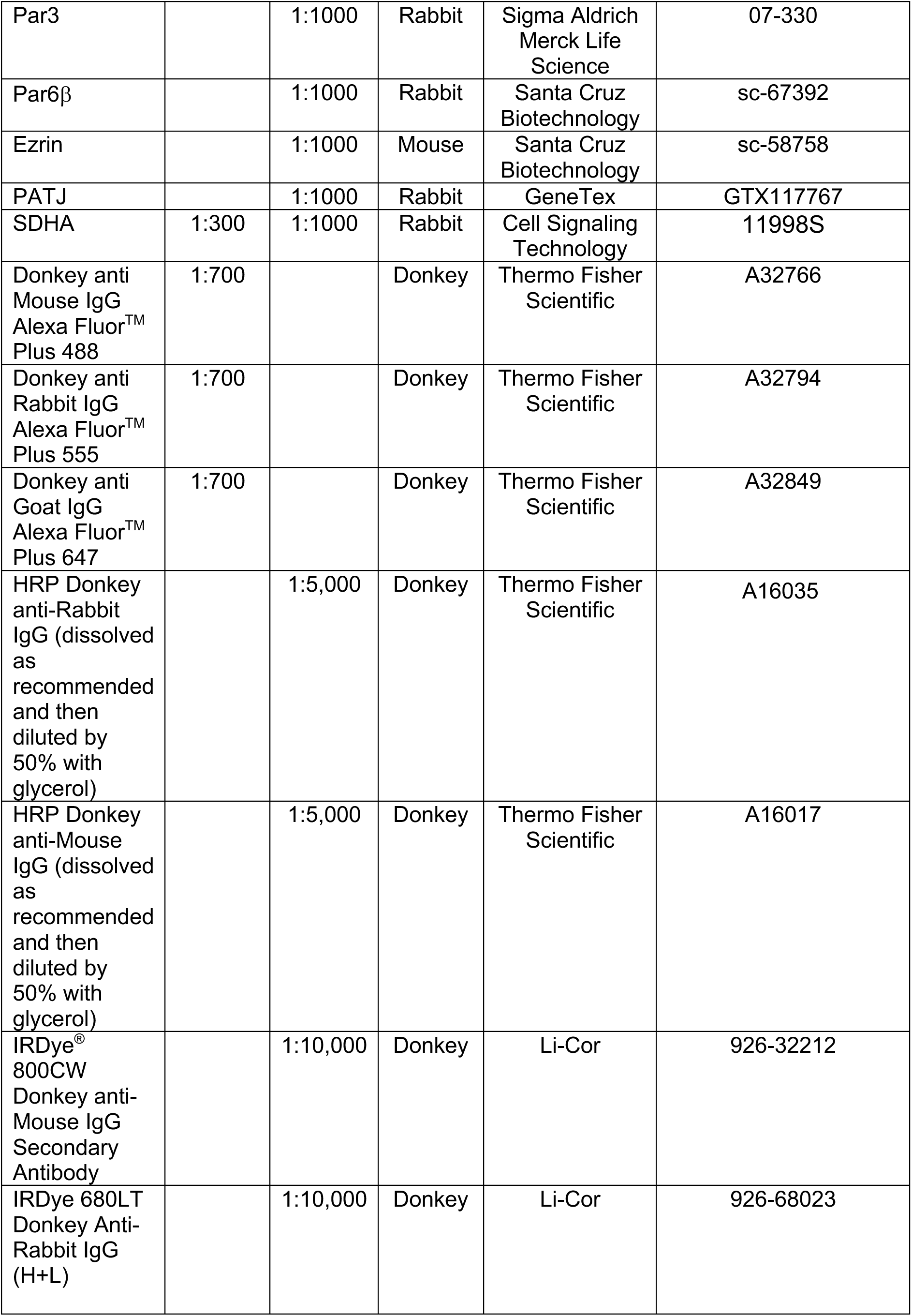
Primary and secondary antibodies used in this study.

| Antibody | Dilution for IHC | Dilution for WB | Species | Company | Catalogue Number |
| --- | --- | --- | --- | --- | --- |
| Arhgef18 | 1:300 | 1:1000 | Rabbit | GeneTex | GTX102223 |
| Arrestin C | 1:300 |  | Rabbit | Proteintech | 11100-2-AP |
| Chx10 (D-11) | 1:300 |  | Mouse | Santa Cruz Biotechnology | Sc-374151 |
| Cre Recombinase (D7L7L) XP® | 1:300 |  | Rabbit | Cell Signaling Technology | 15036S |
| GFAP | 1:300 |  | Rabbit | DAKO | Z0334 |
| Glutamine Synthetase | 1:500 |  | Mouse | BD Biosciences | 610517 |
| Pals1 (G-5) | 1:300 | 1:500 | Mouse | Santa Cruz Biotechnology | sc-365411 |
| p120 Catenin | 1:300 |  | Goat | Santa Cruz Technologies | sc-1730 |
| PKCζ (H-1) | 1:300 | | Mouse | Santa Cruz Biotechnology | sc-17781 |
| pMLC – S19 | 1:300 |  | Mouse | Cell Signaling Technology | 3675S |
| ppMLC – T18,S19 | 1:300 |  | Rabbit | Cell Signaling Technology | 95777S |
| Recoverin | 1:1000 |  | Rabbit | Merck Millipore | AB5585-I |
| Rhodopsin clone 4D2 | 1:1000 |  | Mouse | Merck Millipore | MABN15 |
| Sox9 | 1:300 |  | Rabbit | Merck Millipore | AB5535 |
| Vimentin | 1:300 |  | Mouse | Santa Cruz Biotechnology | sc-373717 |
| ZO-1 (ZO1-1A12) | 1:300 |  | Mouse | Thermo Fisher Scientific | 33-9100 |
| ZO-1 |  | 1:2000 | Rabbit | Matter and Balda | [10.1083/jcb.200211047](https://doi.org/10.1083/jcb.200211047) |
| ZO-1 |  | 1:4000 | Rabbit | Matter and Balda | [10.1083/jcb.200211047](https://doi.org/10.1083/jcb.200211047) |
| N-cadherin (3B9) | 1:100 | 1:500 | Mouse | Thermo Fisher Scientific | 333900 |
| alpha-tubulin (AA4.3) |  | 1/20 | Mouse | DSHB (in-house made supernatant) | AA4.3 |
| β-catenin | 1:300 | 1:1000 | Rabbit | Cell Signaling Technology | 8480S |
| p-Ser552 β-catenin | | 1:1000 | Rabbit | Cell Signaling Technology | 5651S |
| TBK1 |  | 1:1000 | Rabbit | Cell Signaling Technology | 3013S |
| p-Ser172 TBK1 |  | 1:1000 | Rabbit | Cell Signaling Technology | 5651S |
| Pan-cadherin |  | 1:1000 | Rabbit | Sigma Aldrich Merck Life Science | C3678 |
| Par3 |  | 1:1000 | Rabbit | Sigma Aldrich<br>Merck Life<br>Science | 07-330 |
| Par6β | | 1:1000 | Rabbit | Santa Cruz<br>Biotechnology | sc-67392 |
| Ezrin |  | 1:1000 | Mouse | Santa Cruz<br>Biotechnology | sc-58758 |
| PATJ |  | 1:1000 | Rabbit | GeneTex | GTX117767 |
| SDHA | 1:300 | 1:1000 | Rabbit | Cell Signaling<br>Technology | 11998S |
| Donkey anti<br>Mouse IgG<br>Alexa Fluor™<br>Plus 488 | 1:700 |  | Donkey | Thermo Fisher<br>Scientific | A32766 |
| Donkey anti<br>Rabbit IgG<br>Alexa Fluor™<br>Plus 555 | 1:700 |  | Donkey | Thermo Fisher<br>Scientific | A32794 |
| Donkey anti<br>Goat IgG<br>Alexa Fluor™<br>Plus 647 | 1:700 |  | Donkey | Thermo Fisher<br>Scientific | A32849 |
| HRP Donkey<br>anti-Rabbit<br>IgG (dissolved<br>as<br>recommended<br>and then<br>diluted by<br>50% with<br>glycerol) |  | 1:5,000 | Donkey | Thermo Fisher<br>Scientific | A16035 |
| HRP Donkey<br>anti-Mouse<br>IgG (dissolved<br>as<br>recommended<br>and then<br>diluted by<br>50% with<br>glycerol) |  | 1:5,000 | Donkey | Thermo Fisher<br>Scientific | A16017 |
| IRDye®<br>800CW<br>Donkey anti-Mouse IgG<br>Secondary<br>Antibody |  | 1:10,000 | Donkey | Li-Cor | 926-32212 |
| IRDye 680LT<br>Donkey Anti-Rabbit IgG<br>(H+L) |  | 1:10,000 | Donkey | Li-Cor | 926-68023 |

## Notes

### Competing Interest Statement

The authors have declared no competing interest.

