## Supplemental Figures for "*Arhgef18* is a component of the outer limiting membrane required for retinal homeostasis"

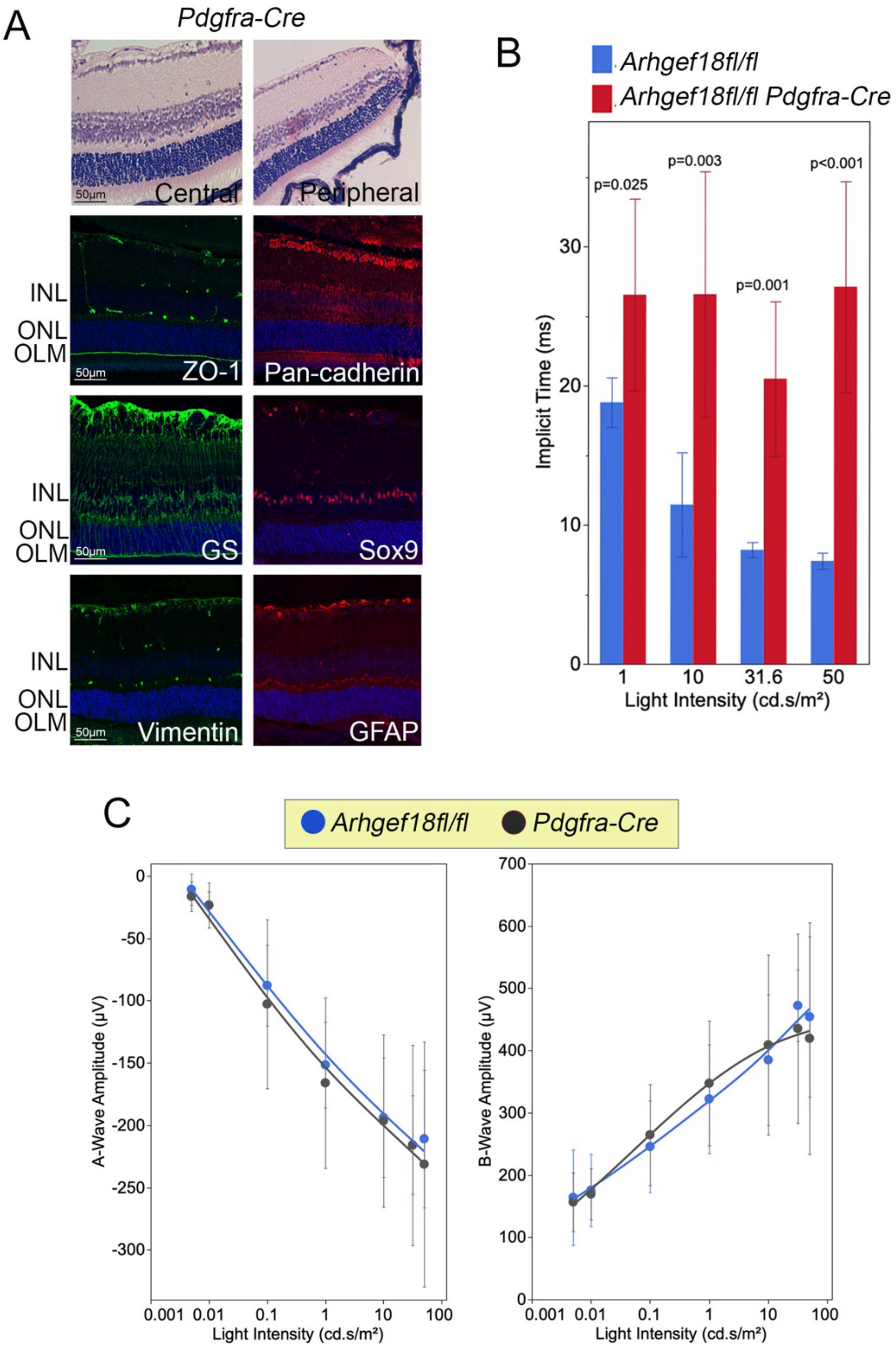


**Figure S1: Analysis of retinal structure and function in C57BL/6J *Pdgfra-Cre mice*.**
(A) Central and peripheral retinas at P21 from C57BL6J *Pdgfra-Cre mice* were stained for H&E (upper panels) or by immunofluorescence for the OLM markers ZO-1 and Pan-Cadherin, the MGC markers GS (glutamine synthetase) and Sox9; and the gliosis markers GFAP (Glial fibrillary acidic protein) and Vimentin. (B) Quantification of A-wave implicit times at different light intensities for *Arhgef18fl/fl* and *Arhgef18fl/fl Pdgfra-Cre* mice (corresponding A-wave amplitudes are shown in figure 1). (*Arhgef18fl/fl*, n=5; *Arhgef18fl/fl Pdgfra-Cre* n=7 animals*).* Shown are means ±1SD; p-values, two-sided t-tests comparing the two genotypes at each light intensity). (C) A and B wave amplitudes at different light intensities of dark-adapted wild-type and *Pdgfra-Cre* ko mice. (*Arhgef18fl/fl*, n=5; *Pdgfra-Cre* n=5 animals*).* Shown are datapoints and fitted curves; p-values, two-sided t-tests comparing the two genotypes at each light intensity).

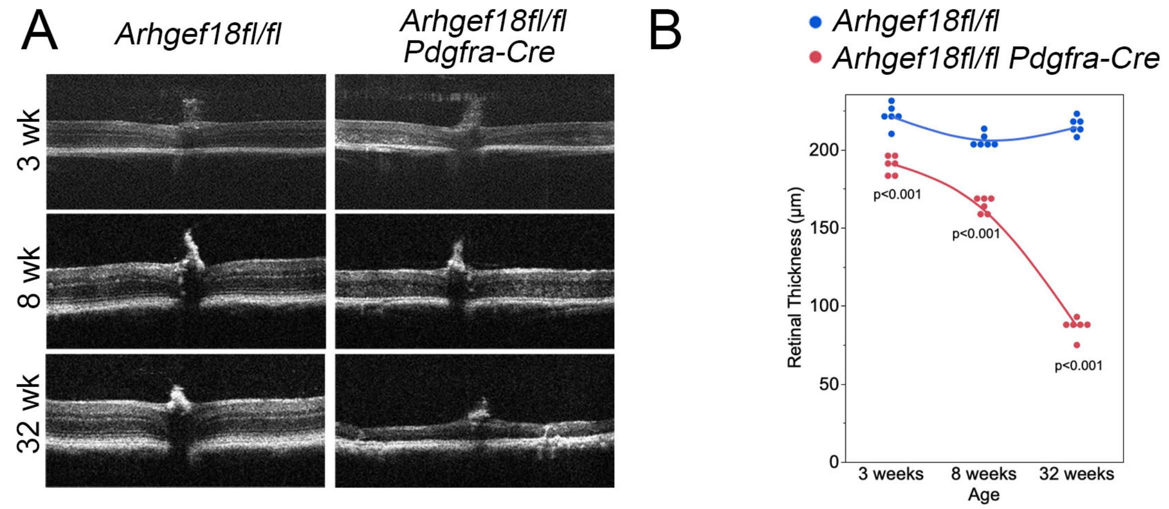


**Figure S2: SD-OCT analysis indicates a progressive retinal degeneration in *Arhgef18fl/fl Pdgfra-Cre mice*.** (A) SD-OCT scans of *Arhgef18fl/fl* and *Arhgef18fl/fl Pdgfra-Cre* mice at 3, 8, and 32 weeks old mice. (B) Quantification of retinal thickness were obtained from the scans (n=6, shown are datapoints and fitted curves, two-sided t-tests comparing the two genotypes at each age).

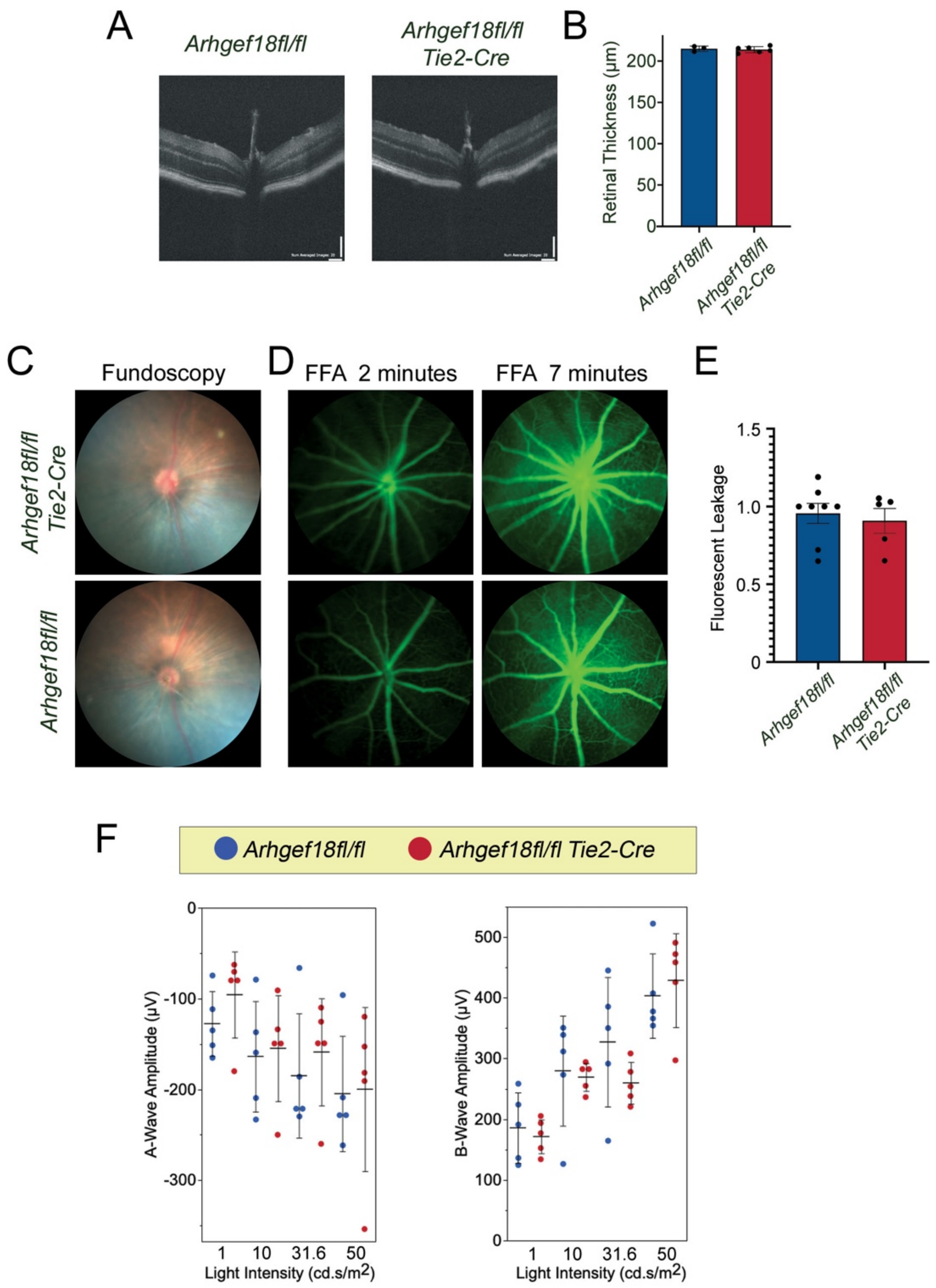


**Figure S3: *Arhgef18fl/fl Tie2-Cre* mice have normal retinal structure and function.**
(A,B) SD-OCT images of *Arhgef18fl/fl* and *Arhgef18fl/fl Tie2-Cre* mice. Panel B shows a quantification of retinal thickness at P21 (shown are datapoints, means ±1SD). (C-E) Fundoscopy (C) and (fluorescent angiography, FFA) of vascular leakage measuring the increase in leakage from 2 to 7 minutes. The quantification shows datapoints and means ±1SD. (F) ERGs at different light intensities of 8-week old *Arhgef18fl/fl* and *Arhgef18fl/fl Tie2-Cre* mice. Shown are A- and B-wave amplitudes of 5 mice for each genotype (means ±1SD; p-values, two-sided t-tests comparing the two genotypes at each light intensity).

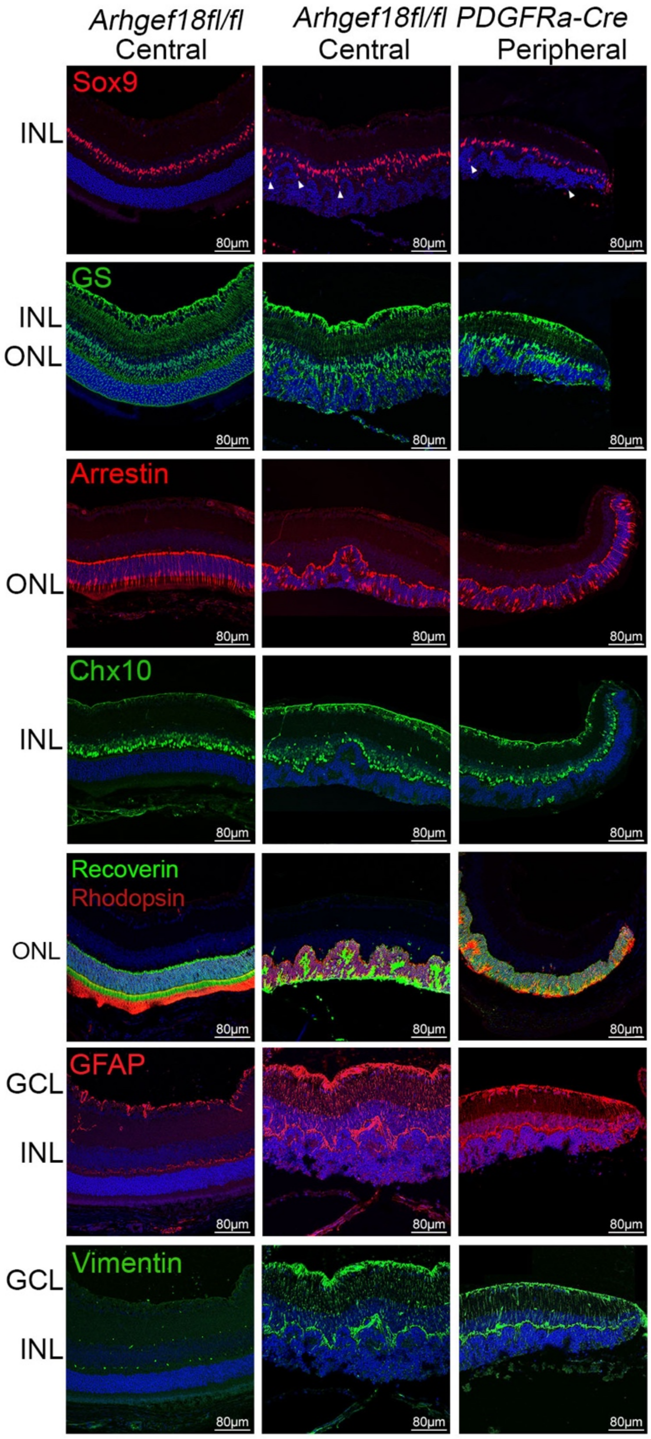


**Figure S4:** ***Arhgef18 Pdgfra-Cre knockout*** **mice display retinal layer disruptions and gliosis.** Immunofluorescence of *Arhgef18fl/fl* and *Arhgef18fl/fl Pdgfra-cre* central and peripheral retinal regions at P21 were stained for retinal markers. Sox9 and glutamine synthetase (GS) for MGs; Chx10 for bipolar cells; and Arrestin, Recoverin and Rhodopsin for photoreceptors. Staining of gliosis markers glial fibrillary acidic protein (GFAP) and vimentin, indicating gliosis across the retina. INL indicates inner nuclear layer, ONL, outer nuclear layer, GCL, Ganglion cell layer.

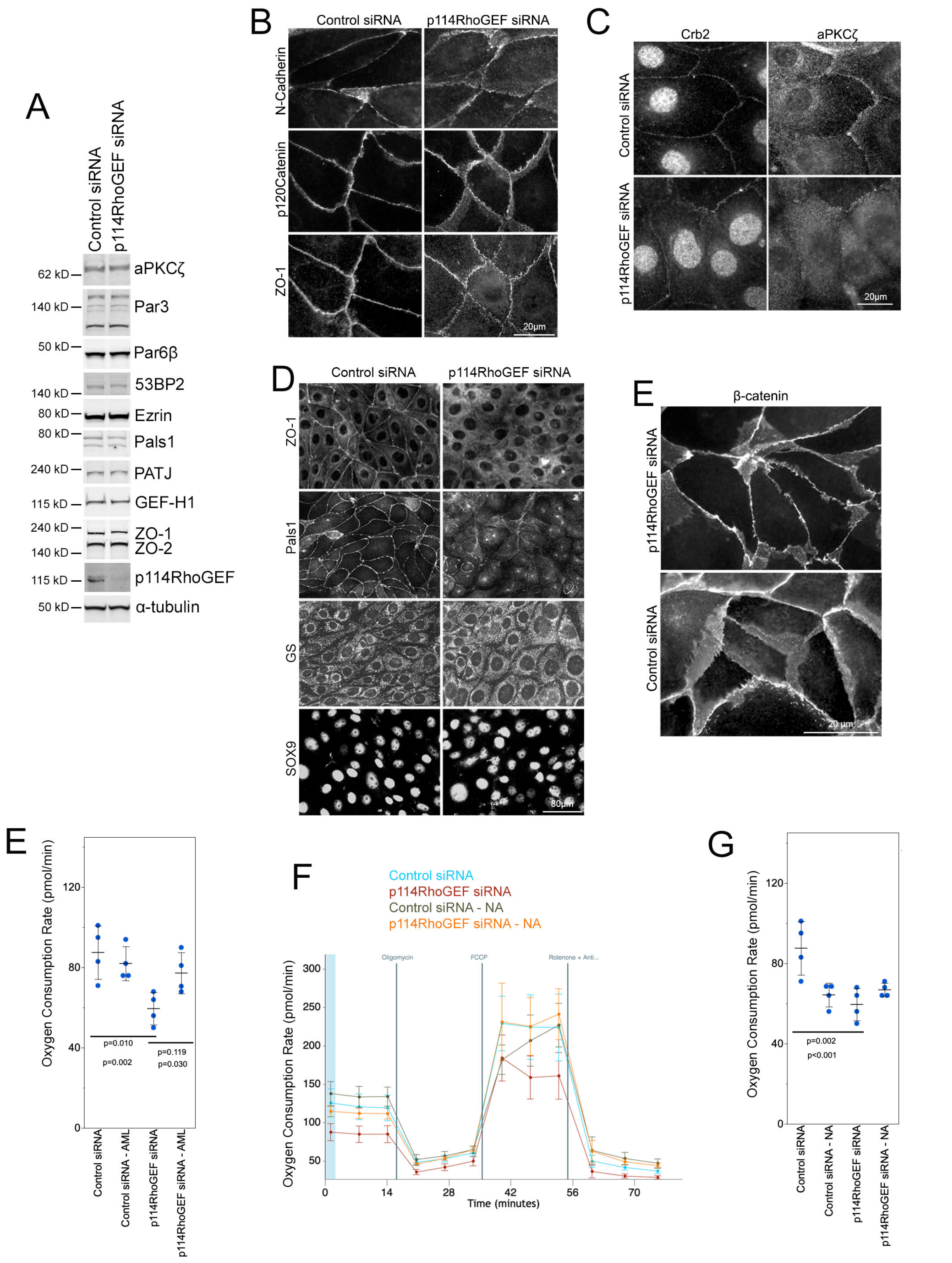


**Figure S5:** (A) Immunoblots and (B-D) Immunofluorescence of the indicated junctional and polarity markers from MIO-M1A cells transfected with nontargeting control or p114RhoGEF siRNAs. The cell extracts in panel A are from the same experiment as figure 6c as the immunoblots are completing the expression analysis. (E) Oxygen consumption rates at basal respiration in siRNA transfected MIO-M1A cells in the absence or presence of Amlexanox (AML). (F-G) Oxygen consumption curves and basal rate calculations in siRNA transfected cells in the absence or presence of Nicotinamide (NA). Panels E and G: n=4; shown are datapoints, means ± 1SD, and p-values from Tukey-Kramer HSD tests.
